# The alpine Edelweiss genome reveals a recent allotetraploid origin and localized subgenome dominance

**DOI:** 10.64898/2026.10.25.761678

**Authors:** Rohan Bhardwaj, Jane Grimwood, Chris Plott, Samija Amar, Flavia Domizia Nardi, Binh Thanh Vo, Jerry Jenkins, Korbinian Schneeberger, Jeremy Schmutz, Hanno Schäfer, Andreas Tribsch, Frank Johannes

**Affiliations:** Plant Epigenomics, Technical University of Munich, 85354 Freising, Germany; HudsonAlpha Institute for Biotechnology, HudsonAlpha Genome Sequencing Center, Huntsville, AL 35806, USA; Department of Genome Plasticity and Computational Genetics, Max Planck Institute for Plant Breeding Research, 50829 Cologne, Germany; Department of Environment and Biodiversity, Universität Salzburg, 5020 Salzburg, Austria; Plant Biodiversity Research, Technical University of Munich, 85354 Freising, Germany; Centre for Research in Agricultural Genomics (CRAG), CSIC-IRTA-UAB-UB, Campus UAB, Bellaterra, Barcelona, Spain; Institute for Crop Biology, Faculty of Mathematics and Natural Sciences, Heinrich-Heine University, Düsseldorf, Germany; Cluster of Excellence on Plant Sciences, Heinrich-Heine University, Düsseldorf, Germany; Faculty of Biology, LMU Munich, Planegg-Martinsried, Germany

**Keywords:** Edelweiss genome, Leontopodium nivale, allotetraploid evolution, subgenome dominance, alpine adaptation, epigenomics

## Abstract

Few plants symbolize the European Alps as profoundly as Edelweiss (*Leontopodium nivale* subsp. *alpinum*), yet its evolutionary origin and genomic constitution has remained elusive. We present a haplotype-resolved, chromosome-scale assembly of an *L. n. alpinum* accession from the Central Alps. The 3.15-Gb genome resolved 48 chromosomes in four phased sets with balanced gene and repeat landscapes. Phylogenomics partitioned these sets into two progenitor-derived subgenomes, DE and FG, demonstrating an allotetraploid origin. Comparison with the diploid subspecies *L. n. nivale* from Southern European Mountains (Appennines) linked DE to this sampled southern lineage, whereas FG derives from another distinct, unresolved diploid progenitor. Chloroplast genomes placed the plastid split between central Alps and Apennine samples in the recent Pleistocene, approximately 20,000 years ago, and LTR insertion-age profiles revealed recent retrotransposon activity consistent with genome shock following allopolyploidization. Although the subgenomes are broadly balanced, large inversions and elevated divergence concentrate on three chromosome groups. On Chromosome 3, structural complexity coincides with FG hypermethylation and DE-biased expression, revealing localized subgenome dominance. Together, these results support previous studies that suggested a westward expansion from the Asian centre of *Leontopodium* diversity, followed by differentiation and hybridization of closely related European lineages. Our assembly positions Edelweiss as an alpine model for recent allotetraploidy and enables genomic studies of demography, selection and high-alpine adaptation.

---

Perhaps no other alpine plant is so deeply embedded in the cultural imagination of the European Alps as the Edelweiss. The distinctive white star of *Leontopodium nivale* subsp*. alpinum* (Cass.) Greuter has become an emblem of Alpine identity, appearing in songs, folk medicine, mountaineering traditions, military insignia and tourist souvenirs [1,2]. Edelweiss grows in exposed, often steep alpine and sunny habitats on dry calcareous grasslands and rocky slopes, mostly around or above 2,000 m [3]. Its felted white bracts and leaves are consistent with adaptation to sunny, windy high-mountain conditions, including protection against ultraviolet radiation [4]. These conspicuous traits and ecological associations have reinforced its image as a flower of alpine landscapes. Although Edelweiss was already recorded in European herbal traditions in the sixteenth century, its modern status as a botanical icon emerged mainly in the nineteenth century, with the rise of alpine tourism and romanticized mountain culture [1,2,5]. As the plant became increasingly sought after by collectors and eager mountaineers, overharvesting prompted early conservation efforts, including legal protection in Austria from 1886 and later protection in Bavaria and Switzerland [2]. Despite this cultural significance and conservation concern, the evolutionary origin and genomic constitution of Edelweiss has remained poorly understood.

The genus *Leontopodium* Cass. (Asteraceae: Gnaphalieae) is predominantly Asian, with its main diversity in Central and Eastern Asian mountains and steppes, especially the Sino-Himalayan region of south-western China, whereas only two closely related Edelweiss taxa occur naturally in Europe [4–6]. Within Europe, *L. nivale* subsp*. alpinum* is broadly distributed across the Pyrenees, Alps, Carpathians and Balkan Peninsula, whereas *L. nivale* subsp*. nivale* Ten. (Tutin) has a restricted and disjunct southern distribution in the central Apennines of Italy, the Pirin Mountains of Bulgaria, and Serbia [4,5,7]. This distribution places central Alpine Edelweiss within a broader Eurasian radiation whose diversity centre lies far to the east [5].

This disjunct east–west Eurasian distribution is consistent with a recent arrival of an Asian lineage into Europe, but molecular studies have not resolved this history in full detail. Nuclear ribosomal and plastid DNA phylogenies recovered *L. n. alpinum* and *L. n. nivale* as a distinct European group within *Leontopodium*, while also showing low nucleotide divergence across the genus and weak resolution among many species [5,6]. AFLP fingerprinting has provided finer geographic structure, separating the European taxa from Mongolian/central Chinese and Yunnan lineages [4]. The same analyses found the highest numbers of private and rare fragments in Yunnan, supporting a longer in situ history in the Sino-Himalayan centre of diversity [4]. More recent plastome and nrDNA analyses have reinforced the view that the *Leontopodium* relationships remain complex, with partly different signals from plastid and nuclear ribosomal data [6]. Together, these studies support the view that the European Edelweiss represents a comparatively young western branch of a recent, Asian-centred radiation. However, they do not explain how the Alpine lineage acquired its present genomic constitution.

Cytological evidence points to aneuploidy and polyploidy as a central mode in speciation. Chromosome-number surveys across *Leontopodium* have suggested several basic chromosome numbers, including *x* = 6, 8, 9 and 11, with deviations from the common *x* = 8 condition occurring toward the edges of the broad Eurasian distribution [8]. European *L. n. alpinum* has been reported with high chromosome numbers, including 2*n* = 48 and 2*n* = 52, consistent with a tetraploid or hexaploid constitution in central European Alpine populations [8]. At the same time, *L. n. nivale* from Gran Sasso (Apennines), Italy, was only recently reported with 2*n* = 24, a count interpreted as tetraploid under *x* = 6 but also compatible with diploidy under *x* = 12 [8]. This ambiguity leaves open the possibility that southern European diploid lineages contributed to the ancestry of the tetraploid *L. n. alpinum* in the central Alps. Whether the Alpine Edelweiss arose through autopolyploid genome duplication or through hybridization of diploid lineages is unknown.

Resolving this distinction matters because auto- and allopolyploid formation can have different consequences for genome evolution. Polyploidy has been a major force in plant diversification, but newly formed polyploids often face challenges of meiotic stabilization, genome reorganization and regulatory conflict [9–12]. In allopolyploids, the merger of related but distinct genomes could trigger a genomic shock, including structural genomic rearrangements, altered transposable element (TE) regulation, epigenetic remodelling and non-additive gene expression [12–15]. Such processes can ultimately lead to subgenome dominance, in which progenitor-derived genomes contribute unequally to gene retention, DNA methylation or expression [13–15]. Studies have shown that such asymmetry can arise rapidly even between very closely related progenitor lineages [16,17].

Here, we begin to address these questions by generating a haplotype-resolved, chromosome-scale genome assembly of an *L. n. alpinum* accession from the Central Alps in Austria and integrating it with transcriptome data, DNA methylation and comparative genomic data from the related Apennine subspecies *L. n. nivale*, sampled in the Gran Sasso region of Italy. We show that the Austrian tetraploid has a recent allotetraploid origin involving two closely related progenitor-derived subgenomes, one of which is linked to the sampled Italian diploid lineage. Although these subgenomes are broadly balanced in gene and repeat content, they differ in chromosome-scale structure, DNA methylation and localized expression patterns, culminating in subgenome dominance on structurally complex Chromosome 3. Beyond resolving the genomic origin of Alpine Edelweiss, the nuclear reference genome provides a framework for population-genomic analyses of migration, introgression and demographic history. Such analyses should help disentangle the complex phylogenetic and phylogeographic relationships among *Leontopodium* species and lineages within Europe and eastward in the Asian centre of diversification We position also Edelweiss as a potential system for studying the recent, and potentially ongoing, genomic consequences of allotetraploidy between closely related lineages.

## Results

### A chromosome-scale assembly and annotation resolve the Edelweiss tetraploid genome

We generated a haplotype-resolved, chromosome-scale assembly of Austrian *Leontopodium nivale* subsp*. alpinum* to establish a genomic framework for testing the origin and organization of the Edelweiss genome (**Online Methods**). Material for genome assembly was collected from a single plant in a natural population in the Zemmgrund, within the Zillertal Alps Nature Park in western Tyrol, Austria, approximately 5 km from the Italian border with South Tyrol/Alto Adige (**Fig. 1a**).

**Fig. 1.**
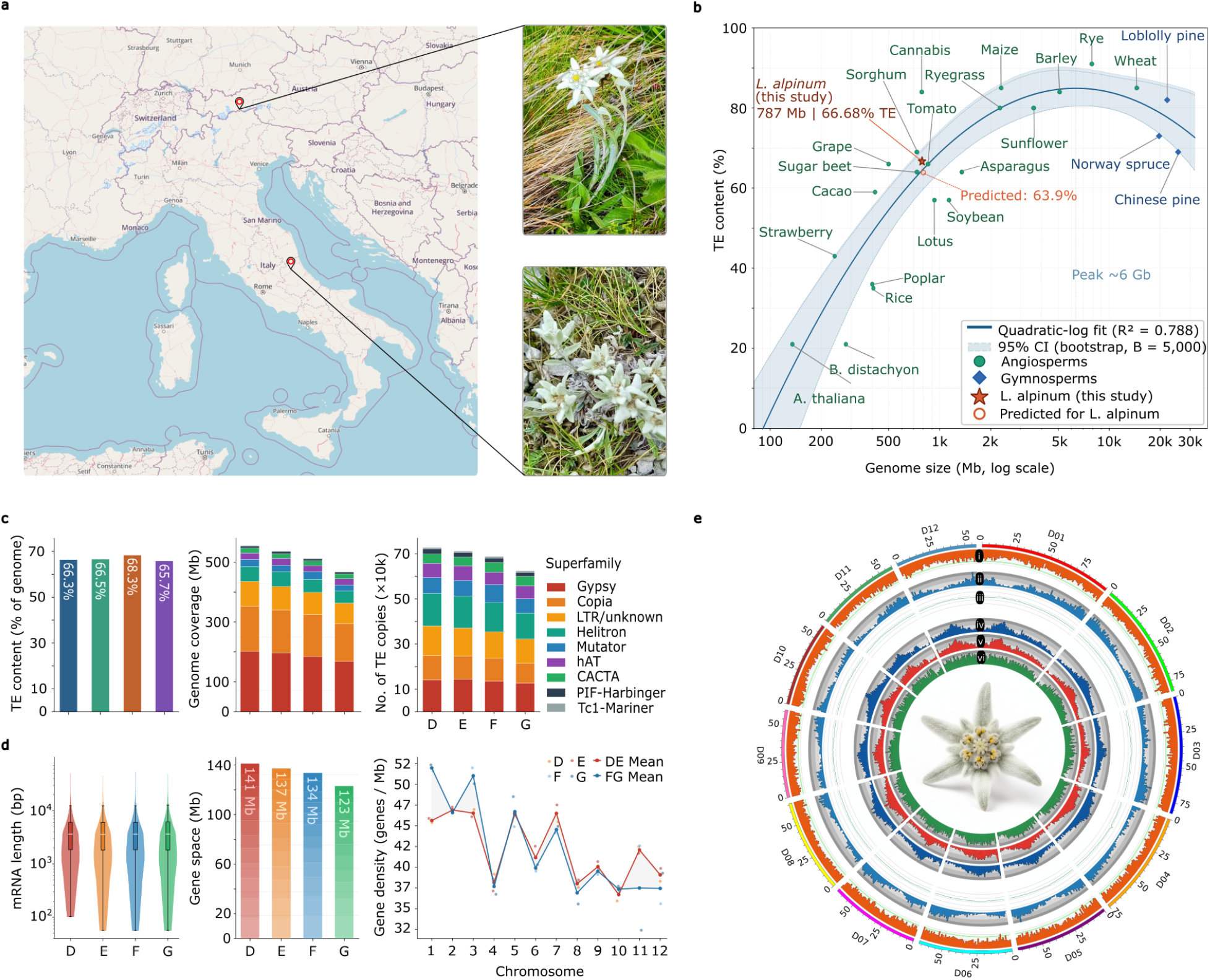
A haplotype-resolved, chromosome-scale assembly resolves four structurally balanced chromosome sets in *Leontopodium nivale* subsp*. alpinum*. **(a)** Sampling localities. The individual used for genome assembly was collected in the Zemmgrund, Zillertal Alps Nature Park, western Tyrol, Austria (∼5 km north of the Austria–Italy border at ∼2,000 m elevation; upper pin). The lower pin marks the *L. n.* subsp. *nivale* locality in Gran Sasso region of the central Apennines, Italy. Inset, *L. n.* subsp. *alpinum* and *L. n.* subsp. *nivale* in its natural alpine habitat. **(b)** Transposable-element (TE) content as a function of genome size for *L. n. alpinum* and 23 chromosome-level angiosperm (green circles) and gymnosperm (blue diamonds) assemblies; the *x*-axis is log-scaled. Solid line, quadratic-log regression (*R*² = 0.788); shaded band, 95% confidence interval from 5,000 bootstrap replicates. The filled red star denotes the observed *L. n. alpinum* value (787 Mb mean chromosome-set size, 66.68% TE); the open orange circle denotes the value predicted from the fit (63.9%), placing the observation 2.8 percentage points above expectation. **(c)** Repeat landscape across the four chromosome sets (D–G). Left, total TE content as a percentage of each set (values inside bars). Middle, genomic coverage (Mb); right, copy number (×10⁴). Middle and right panel’s stacked bars are partitioned by TE superfamily (colour key to the right). **(d)** Gene-space organization across the four chromosome sets. Left, per-set mRNA length distribution (violin, full distribution; box, interquartile range; centre white line, median). Middle, total gene space per set (Mb; values on bars). Right, gene density (genes per Mb) for each of the twelve homoeologous chromosome groups; faint points, individual D, E, F and G values; lines, the mean of the D and E sets (DE, red) and of the F and G sets (FG, blue). **(e)** Circos plot of subgenome D, showing six tracks in 1 Mb bins: (i) TE density (∼65–67%), (ii) gene density, (iii) GC content (∼36%), (iv) CG methylation, (v) CHG methylation, and (vi) CHH methylation. Refer to Supplementary Figure 1 for subgenomes E, F, and G.

The assembly spanned 3.15 Gb across 48 chromosome-scale scaffolds, with a scaffold N50 of 64.8 Mb and a contig N50 of 22.0 Mb (**Table 1**). The recovery of 48 chromosome-scale scaffolds, organized into four sets of 12 chromosomes, is consistent with a tetraploid genome structure. Gaps accounted for only 0.1% of the assembly, indicating high contiguity after long-read assembly and Hi-C scaffolding. The 48 chromosomes were phased into four chromosome sets, designated D, E, F, and G, each containing 12 chromosomes. Chromosome sizes ranged from 44.3 Mb on G11 to 95.2 Mb on D01, and the size hierarchy D > E > F > G was maintained across all chromosome groups (**Supplementary Table 1a,b**). Assembly completeness was high. BUSCO analysis against the embryophyta_odb10 dataset recovered 99.5% of the 1,614 expected single-copy orthologues as complete, and 98.8% of complete BUSCOs were present in duplicated form, consistent with broad representation of the tetraploid gene complement (**Supplementary Table 1c**).

**Table 1.** Genome assembly statistics for the complete tetraploid genome of L. n. alpinum (Austria). Assembly-wide quality metrics (scaffold N50, contig N50) are reported for the full 48-chromosome assembly.

|  |  |
| --- | --- |
| **Total scaffolds** | 48 |
| **Total contigs** | 228 |
| **Assembly size (scaffolds)** | 3,149.0 Mb |
| **Assembly size (contigs)** | 3,147.2 Mb (0.1% gaps) |
| **Scaffold N50 / L50** | 64.8 Mb / 21 |
| **Contig N50 / L50** | 22.0 Mb / 41 |
| **Number of scaffolds >50 kb** | 48 |
| **% genome in scaffolds** | 100% |

We annotated the repeat and gene landscapes of the phased assembly. Repetitive DNA accounted for 65–68% of each chromosome set, with broadly similar repeat profiles across D, E, F, and G. To place this repeat content in a broader plant-genome context, we compared the average *L. n. alpinum* chromosome-set size of 787 Mb and 66.68% transposable-element content with 23 chromosome-level angiosperm and gymnosperm assemblies. Across this comparison, *L. n. alpinum* fell close to the expected relationship between genome size and transposable-element content, with observed transposable-element content only 2.8 percentage points above the predicted value of 63.9% (**Fig. 1b**; **Supplementary Table 2**).

LTR retrotransposons dominated the repeat landscape in all four chromosome sets. Gypsy elements were slightly more abundant than Copia elements, followed by unclassified LTRs, Helitrons, and TIR elements, including Mutator and hAT families (**Fig. 1c**). The highly similar repeat composition across D, E, F, and G indicated that no chromosome set had undergone a major expansion of a specific repeat class. Gene annotation recovered 134,698 protein-coding genes and 165,150 transcripts across the genome, with an average of 1.23 transcripts per gene. Absolute gene number scaled with chromosome-set size: D, the largest set at 841.5 Mb, contained 35,564 protein-coding genes, whereas G, the smallest set at 715.7 Mb, contained 30,663 genes. By contrast, gene density was nearly constant across all four chromosome sets, ranging from 42.26 to 43.15 genes per Mb, and gene space occupied 16.8–17.2% of each set. Mean gene length, mean coding-sequence length, and exon number were likewise similar across chromosome sets (**Fig. 1d**; **Table 2**). Annotation quality was high, with 85% of transcripts supported by RNA-seq data and 95.1% of gene models having an Annotation Edit Distance of ≤0.5. Thus, the phased assembly resolved four balanced chromosome sets with similar repeat composition, gene density, and gene-space organization.

**Table 2.** Annotation statistics for each of the four subgenomes (D, E, F, G) and the complete tetraploid assembly of L. n. alpinum (Austria). Gene density is consistent across subgenomes despite differences in total genome size, indicating proportional gene space partitioning following polyploidization.

|  | **D** | **E** | **F** | **G** | **Whole genome** |
| --- | --- | --- | --- | --- | --- |
| **Chromosomes** | 12 | 12 | 12 | 12 | 48 |
| **Genome size (Mb)** | 841.5 | 811.1 | 780.7 | 715.7 | 3,149.0 |
| **Protein-coding genes** | 35,564 | 34,783 | 33,688 | 30,663 | 134,698 |
| **Transcripts** | 43,666 | 42,529 | 41,280 | 37,675 | 165,150 |
| **Mean gene length (bp)** | 3,970 | 3,946 | 3,970 | 4,014 | 3,974 |
| **Mean CDS length (bp)** | 1,238 | 1,227 | 1,224 | 1,236 | 1,231 |
| **Gene space (Mb)** | 141.2 | 137.3 | 133.8 | 123.1 | 535.3 |
| **Gene space (%)** | 16.78 | 16.92 | 17.13 | 17.20 | 17.00 |
| **Gene density (genes/Mb)** | 42.26 | 42.88 | 43.15 | 42.84 | 42.78 |

### Phylogenomics partitions the tetraploid genome into two subgenomes

Having established a phased tetraploid genome structure, we next asked whether the four chromosome sets reflected duplication within a single lineage or merger between divergent progenitor lineages. Under an autotetraploid origin, the four chromosome sets would not be expected to partition consistently into two genome-wide ancestry groups. By contrast, an allotetraploid origin should retain a phylogenomic signal of two progenitor-derived subgenomes (**Fig. 2a**).

**Fig. 2.**
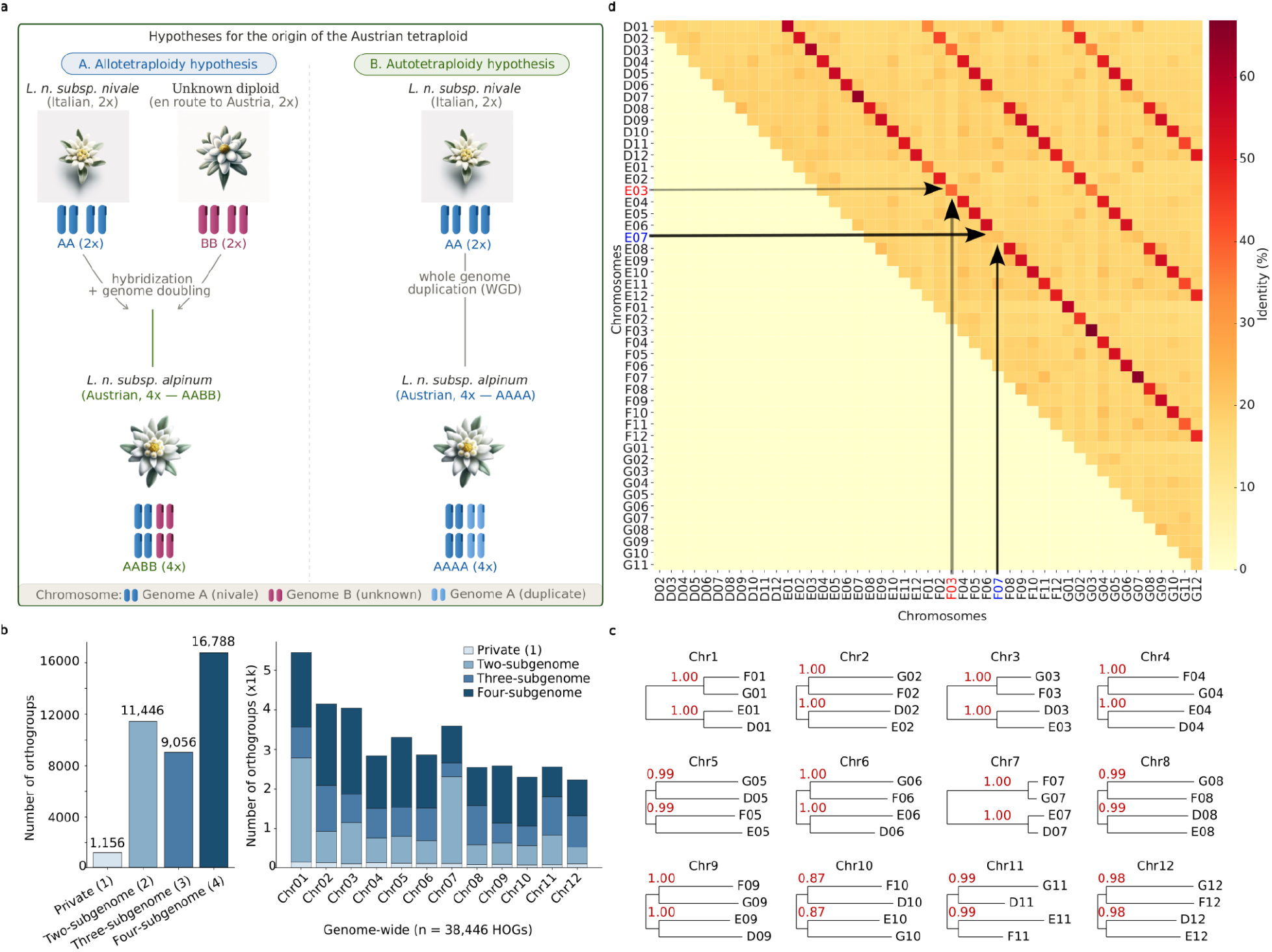
Phylogenomics and pairwise identity partition the tetraploid genome into two closely related subgenomes (DE and FG), supporting an allotetraploid origin. **(a)** Competing hypotheses for the origin of the Austrian tetraploid. Under allotetraploidy (left), hybridization between *L. n.* subsp. *nivale* (Italian, 2x; genome A) and an unknown diploid lineage (genome B), followed by genome doubling, yields an AABB tetraploid. Under autotetraploidy (right), whole-genome duplication of a single *L. n. nivale*-like lineage yields an AAAA tetraploid. Chromosome cartoons are colored by ancestry (key, bottom): genome A (*L. n. nivale*), genome B (unknown), and the duplicated genome-A copy. **(b)** Orthogroup representation across the four chromosome sets. Left, genome-wide counts of hierarchical orthogroups (HOGs) partitioned by the number of chromosome sets in which they are represented, private to one set, or shared across two, three or four sets (values above bars; *n* = 38,446 HOGs). Right, the same four categories resolved per homoeologous chromosome group (Chr01–Chr12); stack shading as in the key. **(c)** Rooted species trees for each of the 12 homoeologous chromosome groups, inferred with STAG from single-copy orthogroups. Each tree relates the four phased copies (D, E, F, G) of one chromosome group. Red values are STAG support. All chromosomes recover two groupings, indicating separate lineages. **(d)** Pairwise sequence identity between all 48 phased chromosomes (D01–G12), colored by percent identity (key, right). The strong diagonal marks each chromosome against its homoeologues. Arrows and highlighted axis labels mark representative cases for chromosome groups 03 and 07: the between-subgenome E–F comparison shows markedly reduced identity, whereas the corresponding within-subgenome pairs (D07–E07 and F07–G07) retain high identity.

We first examined orthogroup representation across the four chromosome sets (**Fig. 2b**). Genome-wide, orthogroups represented in all four chromosome sets formed the largest class, comprising 16,788 of 38,446 hierarchical orthogroups (43.7%). Orthogroups represented in two chromosome sets accounted for 11,446 cases (29.8%), and three chromosome sets accounted for 9,056 cases (23.6%). Private orthogroups restricted to a single chromosome set were rare, comprising 1,156 cases (3.0%; **Supplementary Table 3a**). Thus, a large fraction of the gene complement was retained across all four chromosome sets, providing a suitable basis for testing their evolutionary relationships.

We inferred phylogenetic relationships among the four chromosome sets using 9,961 conserved orthogroups represented by a single copy in each set. These analyses recovered a consistent partition in which D clustered with E, and F clustered with G, across 9 of 12 chromosome groups (**Fig. 2c**). Orthogroup-based clustering resolved the four subgenomes into two lineage pairs. This repeated grouping supports an allotetraploid origin in which the tetraploid genome derives from two progenitor lineages. Pairwise whole-genome sequence identity supported the same partition. Within-pair comparisons between D and E, and between F and G, showed uniformly higher identity across chromosome groups than comparisons between the DE and FG pairs (**Fig. 2d**).

### The DE subgenome is linked to a southern European diploid lineage

To begin to understand the evolutionary origin of the two subgenomes, we assessed whether either subgenome could be linked to the southern European lineage highlighted by previous cytological work. Stille et al. reported a chromosome count of 2*n* = 24 for *L. nivale* from Gran Sasso, Italy, and noted that this count could be interpreted either as tetraploidy under *x* = 6 or as diploidy under *x* = 12 [8]. We therefore collected material from single plants in the Gran Sasso region (prot. 0009734/2023) corresponding to the locality reported by Stille et al. and generated Illumina short-read sequence data from Italian *L. n. nivale* (**Online Methods; Fig. 1a**).

Using k-mer-based estimates [18,19], we found that the Italian *L. n. nivale* sample had a haploid genome size of approximately 1.14 Gb and heterozygosity of 2.14%. Its k-mer frequency profile was best fit by a diploid model, with a model fit of 93.2% (**Fig. 3a**). The elevated signal at approximately 150× coverage was attributable to collapsed repetitive sequences rather than to an additional ploidy peak (**Fig. 3a**). These results support the diploid interpretation of the 2*n* = 24 chromosome count reported for *L. n. nivale* from Gran Sasso. The Italian sample therefore provided a suitable lineage for testing whether one of the two Austrian subgenomes shows greater affinity to the southern European *L. n. nivale* lineage.

**Fig. 3.**
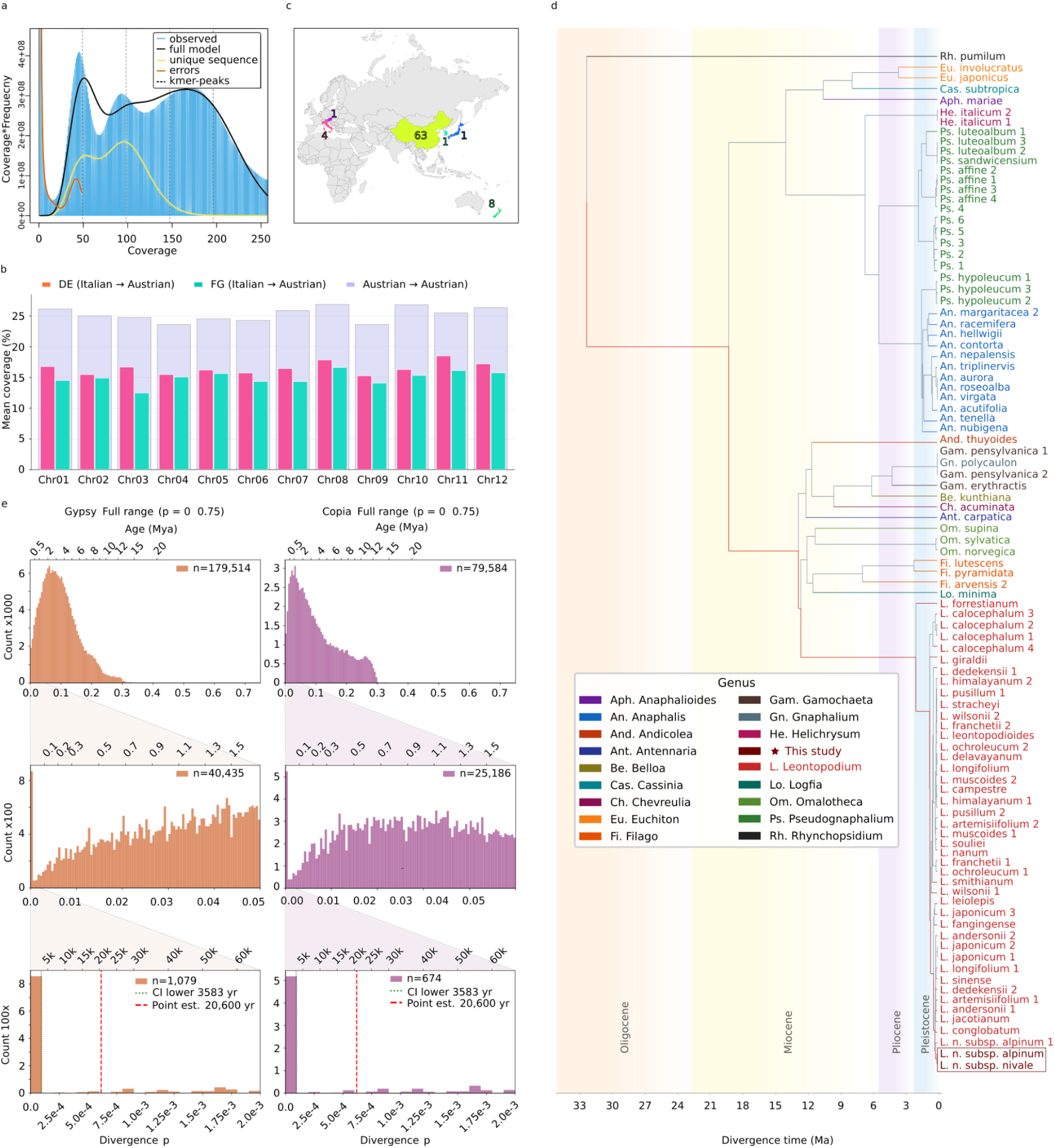
The DE subgenome shows greatest affinity to a southern European diploid lineage, and chloroplast dating places the Austrian–Italian divergence in the Late Pleistocene. **(a)** *k*-mer frequency spectrum of Italian *L. n.* subsp. *nivale* (Gran Sasso) generated using GenomeScope2.0. The profile is best explained by a diploid model (model fit 93.2%; haploid genome size ≈ 1.14 Gb; heterozygosity 2.14%). The elevated signal at approximately 150× coverage reflects collapsed repetitive sequence rather than an additional ploidy peak, consistent with the diploid interpretation of the reported 2*n* = 24 count. **(b)** Mean short-read coverage of Italian *L. n. nivale* reads aligned to the Austrian *L. n. alpinum* assembly (MAPQ ≥ 30) for each homoeologous chromosome group. Within each group, the orthogroup-defined homoeologue pair with higher Italian-read coverage was designated DE (pink) and the other pair FG (green). This required relabelling the original size-based names in 5 of the 12 groups (Chr04, Chr05, Chr07, Chr10 and Chr11). Coloured bases show Italian-read coverage over DE and FG. The purple cap shows coverage of Austrian reads self-aligned to the same assembly under identical filtering (≈25%), which serves as the mappability baseline. Mean genome-wide Italian-read coverage was 16.48% over DE and 14.92% over FG. **(c)** Geographic origin of the sampled accessions included in the chloroplast phylogeny, coloured by region; numbers give the count of accessions per region. Precise locations of 22 samples were not provided in their original submissions. **(d)** Time-calibrated chloroplast phylogeny of 96 Asteraceae taxa, including 40 *Leontopodium* accessions and three European accessions (the two samples of this study marked by a star). Tips are coloured by genus (key); numbers at nodes are median divergence-time estimates (Ma); background shading marks geological epochs. Austrian *L. n. alpinum* and Italian *L. n. nivale* are recovered as sister lineages (red branch), whereas the sampled Asian *Leontopodium* lineages diverge substantially earlier. RelTime placed the Austrian–Italian split at ≈ 20,600 years ago (95% CI 3,583–118,599 years). **(e)** LTR-retrotransposon insertion-age profiles for *Gypsy* (orange, left) and *Copia* (purple, right) elements across the four chromosome sets, shown at three successive zoom levels of sequence divergence (*p*, bottom axis; corresponding age in Mya, top axis; *n* per panel inset). Top, full range (*p* = 0–0.75): *Gypsy* peaks at *p* ≈ 0.08–0.12 (≈ 2–4 Mya) while *Copia* is younger overall. Middle, *p* = 0–0.05. Bottom, *p* = 0–0.002: both families show an excess of near-zero-divergence insertions; the chloroplast point estimate (20,600 years, red dashed) and the lower confidence bound (3,853 years, green dotted) are marked for reference.

To test this, we aligned short-read sequences from Italian *L. n. nivale* to the Austrian *L. n. alpinum* assembly and compared coverage (MAPQ ≥ 30) between the two orthogroup-defined homoeologue pairs within each of the 12 chromosome groups (**Fig. 3b; Supplementary Table 4**). In every group, both homoeologues of one clustered pair showed higher Italian coverage than both homoeologues of the other, so the homoeologues that cluster together also share the same ancestral affinity. As an alternative approach, we aligned Italian *L. n. nivale* reads separately to each Austrian subgenome and called variants in single-copy orthologous coding sequences; the Italian samples showed small but fixed differences against DE and FG (1.88 vs. 1.91 homozygous-alternative sites per kb) (**Supplementary Table 4**). We therefore designated the pair with greater affinity to the Italian lineage DE and the other FG. This required relabelling the original size-based chromosome names in 5 of the 12 groups (Chr04, Chr05, Chr07, Chr10 and Chr11), whereas in the remaining 7 groups the original labels already matched this grouping. All analyses below use this lineage-based DE/FG nomenclature. The overall divergence between DE and FG was modest: mean coverage was 16.48% across DE and 14.92% across FG, indicating that the two progenitor-derived lineages are closely related. Nevertheless, the genome-wide consistency of the DE/FG split showed that the four chromosome sets were not equivalent copies of a single ancestral genome.

To place the absolute coverage values in context, we also aligned short-read data from the Austrian reference individual back to its own assembly using the same filtering criteria (**Fig. 3b**). This self-alignment recovered approximately 25% coverage. Thus, although Italian reads mapped preferentially to DE over FG, their absolute coverage was lower than the Austrian self-alignment baseline. This difference could reflect divergence between the sampled Italian lineage and the exact DE progenitor lineage. It could also reflect structural or repeat changes that accumulated in the Austrian tetraploid after allopolyploidization and reduced the mappability of Italian reads to the present-day assembly. The coverage data therefore identify the sampled Italian *L. n. nivale* lineage as the closest sampled relative of DE, while leaving open whether it represents the immediate DE progenitor or a surviving relative of that lineage.

### Chloroplast divergence and LTR profiles support a recent Pleistocene timescale

To place the relationship between the Austrian tetraploid and the sampled Italian diploid lineage in a temporal framework, we assembled complete chloroplast genomes from Illumina short-read data (**Online Methods**). We integrated these assemblies into a time-calibrated chloroplast phylogeny of 96 Asteraceae taxa, including 40 *Leontopodium* taxa sampled across the China-centred distribution of the genus and three European accessions, including the samples from this study (**Fig. 3c**). Austrian *L. n. alpinum* and Italian *L. n. nivale* formed sister lineages within the *Leontopodium* clade, whereas the sampled Asian *Leontopodium* lineages diverged substantially earlier (**Fig. 3d; Supplementary Table 5**). The two European chloroplast assemblies differed by only approximately 3 SNPs, confirming very shallow plastid divergence.

RelTime analysis placed the Austrian–Italian chloroplast split at approximately 20,600 years ago, with a 95% confidence interval spanning 3,583 to 118,599 years (**Online Methods**). The point estimate falls within the Late Pleistocene and is consistent with earlier biogeographic hypotheses that European *Leontopodium* lineages were shaped by Pleistocene range shifts [2]. Although the confidence interval is broad, this uncertainty is expected for very recently diverged plastid lineages. Because chloroplasts only trace cytoplasmic inheritance, this estimate does not directly date the nuclear genome merger. However, under the simplest scenario in which the Austrian plastid was inherited during allopolyploid formation, the Austrian–Italian plastid split provides an approximate upper bound on the age of tetraploidization.

Polyploidization, especially when coupled with hybridization, can perturb TE regulation and has been associated with rapid TE mobilization or structural and epigenetic reorganization near TE insertion sites in several plant systems [20–23]. We therefore asked whether the Edelweiss genome retained an LTR-retrotransposon signal compatible with recent genome perturbation. We analysed insertion-age profiles of Gypsy and Copia LTR retrotransposons across the four chromosome sets (**Online Methods**). Full-range divergence distributions revealed distinct age profiles for the two element families (**Fig. 3e**). Gypsy elements showed a broad peak at divergence values of approximately 0.08–0.12, corresponding to an estimated age of approximately 2–4 Mya. Copia elements had a younger overall age distribution, with most insertions concentrated at lower divergence values corresponding to approximately 0.5–1 Mya. These profiles indicate that the repeat landscape of Austrian *L. n. alpinum* reflects a longer history of transposon activity predating the shallow Austrian–Italian chloroplast divergence.

We then focused on the youngest end of the LTR divergence distribution. Insertions occurring near the chloroplast point estimate of 20,600 years would be expected to have extremely low divergence, approximately p = 3.0 × 10⁻⁴, below the resolution at which individual insertions can be dated reliably. Indeed, we identified an excess of Gypsy and Copia elements with divergence values of effectively zero, consistent with very recent LTR-retrotransposon activity. Although such undiverged elements cannot refine the chloroplast-based estimate or be assigned to a specific historical event, their presence is consistent with recent retrotransposon mobilization, as expected under genome perturbation following allopolyploidization.

### Chromosome-scale rearrangements concentrate on three homoeologous chromosome groups

Having established a recent allotetraploid origin, we examined how the two progenitor-derived nuclear subgenomes are organized at chromosome scale in more detail. This was important because the structural similarity of homoeologous chromosomes can influence meiotic behavior in polyploids. Highly similar chromosome sets may retain the potential for homoeologous pairing and exchange, whereas increased sequence and structural divergence can contribute to diploid-like meiotic behavior by limiting recombination between subgenomes [24,25]. Large inversions are particularly relevant in this context because they can suppress crossing over between alternative chromosomal arrangements and thereby promote more independent evolution of the affected haplotypes or chromosome segments [26,27]. We therefore compared synteny, inversion structure and nucleotide divergence within and between the DE and FG subgenomes in detail.

Synteny was largely preserved within each subgenome. D–E and F–G comparisons recovered 577 Mb and 493 Mb of collinear sequence, respectively, with total inversion spans of only approximately 25-27 Mb in each comparison (**Fig. 4a**, **Supplementary Table 6a**). By contrast, the inter-subgenome E–F comparison showed reduced collinearity, with 362 Mb of syntenic sequence and large inversions spanning approximately 123 Mb in total. These inversions were not distributed evenly across the genome, but were concentrated on three homoeologous chromosome groups: Chr07, Chr03 and Chr01 (**Fig. 4a**; **Supplementary Table 6b**). Chr07 carried the largest inter-subgenome inversion, spanning approximately 41.8 Mb, followed by Chr03 at approximately 35 Mb and Chr01 at approximately 26.4 Mb. Within-subgenome inversions on the same chromosome groups were much smaller, rarely exceeding 7–10 Mb, indicating that the largest rearrangements distinguished DE from FG rather than homologous chromosome sets within each subgenome (**Fig. 4b**).

**Fig. 4.**
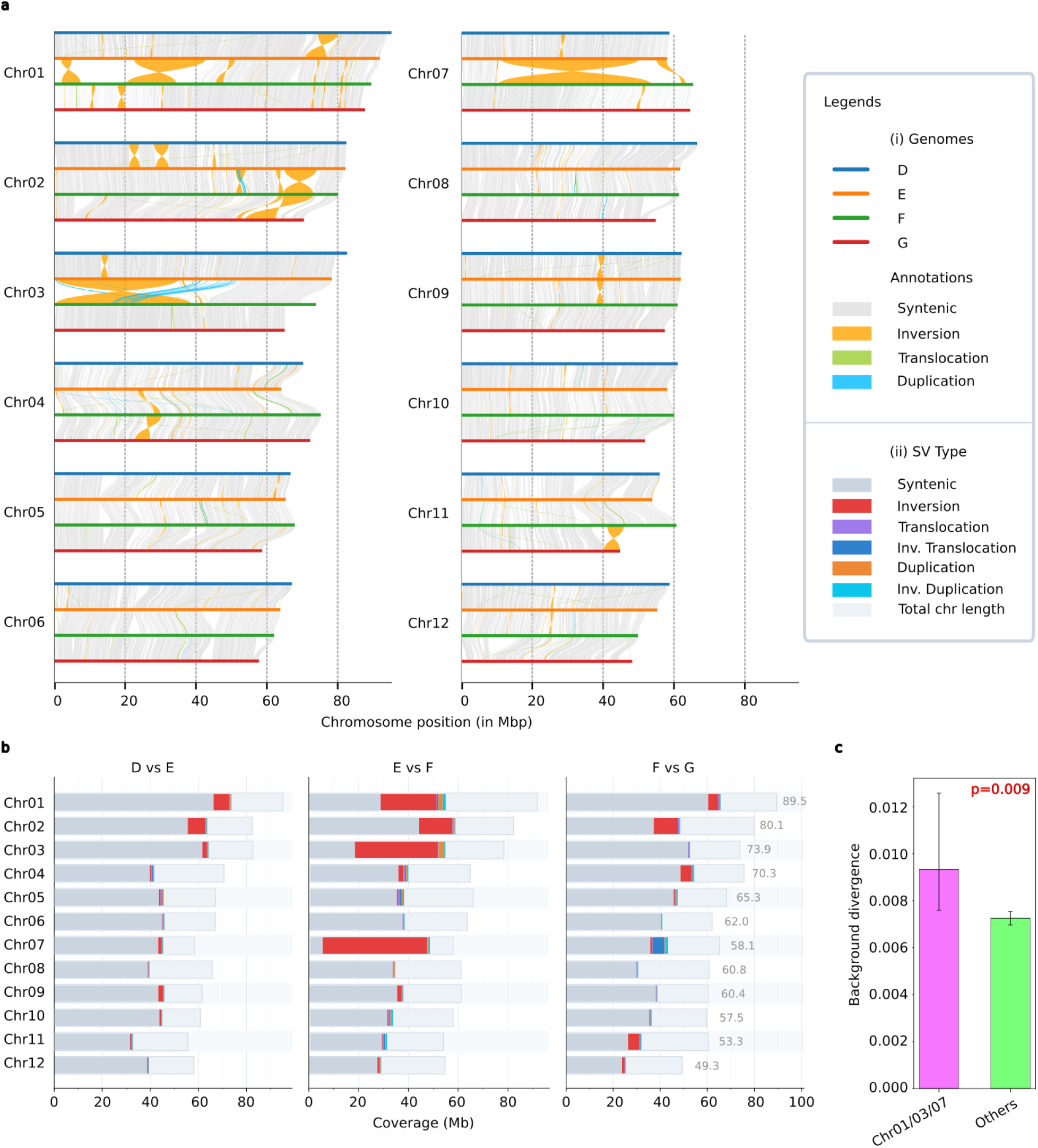
Inter-subgenome structural divergence concentrates on three homoeologous groups, with a nested inversion–duplication architecture on Chr03. **(a)** Pairwise synteny across the four phased copies of each homoeologous chromosome group (Chr01–Chr12). Within each group, the four ideograms are ordered D, E, F, G from top to bottom (genome key, i); ribbons connect vertically adjacent copies and are coloured by relationship (Annotations key): syntenic (grey), inverted (orange), translocated (green) and duplicated (cyan). Large inversions occur predominantly at the inter-subgenome E–F interface and are concentrated on Chr07 (≈ 41.8 Mb), Chr03 (≈ 35 Mb) and Chr01 (≈ 26.4 Mb). On Chr03, duplicated blocks (cyan) lie within the inverted interval, producing a nested inversion–duplication. **(b)** Genomic span (Mb) assigned to each structural class for the three pairwise comparisons within-subgenome D–E (left) and F–G (right), and inter-subgenome E–F (middle), resolved per chromosome group. Segments are coloured by structural-variant type (SV Type key, ii); the faint background bar denotes total chromosome length. Inversion span (red) is far larger in the E–F comparison than in either within-subgenome comparison, and within E–F is greatest on Chr07, Chr03 and Chr01, accompanied by nested inverted-duplication sequence (orange) on Chr03 and Chr01. Friedman omnibus test across the three comparisons (statistic = 12.5, *P* = 0.002). **(c)** Background nucleotide divergence between the DE and FG subgenomes, measured in syntenic sequence outside structural variants and computed from the E–F comparison as a proxy for DE–FG. Per chromosome group, divergence is the number of background SNPs (SNPs residing in syntenic parent blocks) divided by the base-pair-resolved aligned syntenic footprint (SYNAL blocks). Bars, group mean for the three structurally divergent groups (Chr01, Chr03, Chr07; *n* = 3; pink) versus the remaining nine groups (*n* = 9; green); error bars span the within-group minimum-to-maximum range. Background divergence is elevated on the structurally divergent groups, with all three exceeding every other group (*P* = 0.009, two-sided Mann–Whitney U, n = 3 vs 9).

Interestingly, the chromosome groups carrying the largest inter-subgenome inversions also showed elevated DE–FG nucleotide divergence outside the inverted intervals (**Fig. 4c**). This pattern suggests that structural divergence on these chromosomes is associated with broader subgenome differentiation, consistent with reduced recombination or more independent evolutionary histories of the DE and FG chromosomes in these regions. The same chromosome groups also showed distinct orthogroup copy-number patterns. Chr07 showed the strongest departure from the genome-wide four-set orthogroup pattern, with 2,215 orthogroups represented in two chromosome sets compared with only 937 represented in four or more sets (**Fig. 2b**). Chr01 showed a similar, though weaker, shift, with 2,641 two-set orthogroups compared with 1,883 orthogroups represented in four or more sets (**Supplementary Table 3b**). These patterns point to substantial gene loss, disruption or copy-number reorganization on Chr01 and Chr07.

We asked whether genes retained or depleted on these structurally divergent chromosomes showed functional biases. Dosage-balance models predict that genes involved in regulatory networks or stoichiometrically constrained cellular complexes are preferentially retained after polyploidy, whereas other genes more readily return to lower copy number [28,29]. Consistent with this expectation, GO enrichment analysis of retained orthogroups on Chr01 and Chr07 revealed enrichment for functions related to transcriptional regulation, RNA processing, spliceosomal complexes, nuclear pore organization and nucleocytoplasmic transport (**Supplementary Table 7a–f**). These categories overlap with functional classes previously associated with dosage-sensitive gene retention after whole-genome duplication [29]. By contrast, two-copy orthogroups were enriched for a more heterogeneous set of terms (**Supplementary Table 7a–f**). Thus, copy-number reorganization on Chr01 and Chr07 was not functionally random, but was associated with preferential retention of genes involved in nuclear regulatory and RNA-processing machinery. Chr03 showed a different pattern. Although it carried a large inter-subgenome inversion, it retained a majority of four-set orthogroups, with 2,166 orthogroups represented in four or more sets compared with 1,052 represented in two sets. Chr03 therefore combined strong structural divergence with broader preservation of orthologous gene content across the four chromosome sets. This distinction suggested that Chr03 might differ from Chr01 and Chr07 not only in structural organization, but also in the extent to which gene copy number was disrupted.

Duplicated blocks further distinguished the structurally complex chromosome groups. Between the E and F chromosome sets, prominent duplicated segments occurred on Chr03, spanning approximately 1.82 Mb (**Fig. 4b**). Duplicated blocks on all other chromosome groups were smaller than 0.6 Mb. On Chr03, the duplicated blocks overlapped the large inter-subgenome inversion, producing a nested rearrangement in which duplicated sequences lay within a large inverted chromosomal region (**Fig. 4a**). Notably, 132 single-copy orthogroups were retained within these duplicated and inverted Chr03 blocks, indicating that this region preserved orthologous gene content despite local structural complexity.

Together, these analyses show that chromosome-scale divergence between DE and FG is concentrated in a small number of homoeologous groups, with Chr01 and Chr07 displaying pronounced copy-number reorganization, whereas Chr03 preserves much of its orthologous gene content within a nested inversion–duplication architecture.

### Localized subgenome dominance is concentrated on structurally complex Chr03

Structural rearrangements, transposable elements, and hybridization can perturb epigenomic regulation in plant genomes, with potential consequences for gene expression [13–15,20]. We therefore asked whether the structural divergence between the DE and FG subgenomes was accompanied by DNA methylation and transcriptional divergence. To this end, we generated whole-genome bisulfite sequencing data from individuals of the Austrian population used for the reference assembly, together with transcriptome data from petal and rosette tissue (**Online Methods**).

At the genome-wide scale, DE and FG showed a subtle but consistent methylation asymmetry. Overall, FG displayed very similar methylation levels as DE in all three cytosine contexts (meanDE − meanFG, n = 2) by −0.95 pp (percentage points) in CG, −0.89 pp in CHG and −0.26 pp in CHH (**Fig. 5a**). However, because genome-wide methylation averages can mask feature-specific differences, especially when effects occur in opposite directions, we analysed methylation separately across genes and LTR retrotransposons. Across genes, we found significant FG hypermethylation in the CG context across gene bodies and all surrounding regions, whereas CHG and CHH differences extended more broadly across gene bodies and downstream regions (**Fig. 5b**). LTR retrotransposons showed the opposite pattern: within LTR bodies, DE was more highly methylated than FG, particularly in the CHG context (**Fig. 5b**). Thus, DNA methylation differences between the subgenomes were feature-specific, with stronger gene-associated methylation on FG and stronger LTR-body methylation on DE.

**Fig. 5.**
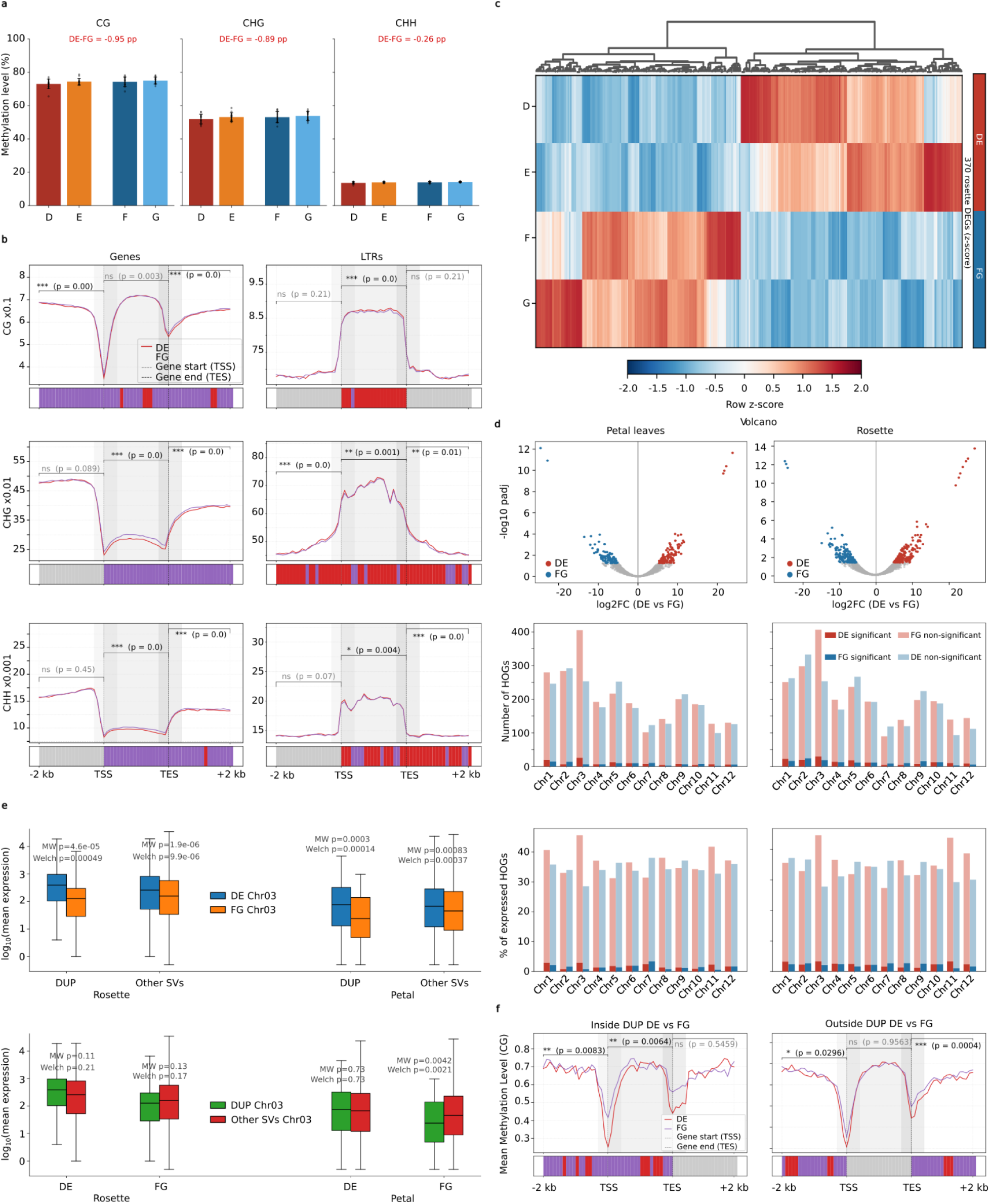
Regulatory divergence between the DE and FG subgenomes is buffered genome-wide but becomes locally pronounced on structurally complex Chr03, where DE-biased expression coincides with FG hypermethylation. **(a)** Genome-wide DNA methylation in the CG, CHG and CHH contexts, compared between subgenomes. Within each context, bars are the mean per-chromosome methylation of the four phased subgenomes (D, E, F, G; warm shades, DE; cool shades, FG); error bars, s.d. across the 12 chromosomes; overlaid points, individual chromosomes. Brackets group D+E (DE) and F+G (FG). pp, difference in methylation level between DE and FG lineages, in percentage points. FG is marginally hypermethylated in all three contexts, with differences below 0.89 pp. **(b)** Metagene methylation profiles across all genes (“Genome-Wide”, left) and LTR retrotransposons (right) for DE (red) and FG (purple), in CG (top), CHG (middle) and CHH (bottom). Each profile spans 2 kb upstream of the transcription start site (TSS), the body (TSS–TES) and 2 kb downstream. Significance is reported per region (upstream, body, downstream) by paired Wilcoxon signed-rank over the bins of that region; the strip beneath each profile marks per-bin DE-versus-FG differences; red, DE higher; purple, FG higher; grey, non-significant (ns). Genes show FG-biased methylation near across all contexts, whereas LTR bodies are highly methylated on DE, in all contexts. **(c)** Expression heatmap of the 370 rosette genes with significant DE-versus-FG bias (row *z*-score across the four subgenomes; columns hierarchically clustered). Rows are the D, E, F and G copies, grouped into DE and FG (annotation on the right). Biased genes partition into DE-high and FG-high sets rather than a uniform genome-wide skew. **(d)** Volcano plots of DE-versus-FG differential expression for petal (left) and rosette (right); *x*, log₂ fold change (DE versus FG); *y*, −log₁₀ adjusted *P*. Significantly DE-biased (red) and FG-biased (blue) genes are highlighted. Below, per-chromosome counts (number of HOGs) and proportion (% of expressed HOGs) of DE-significant, FG-significant and non-significant genes, for petal and rosette. Significant bias affects only a small number of genes on every chromosome. **(e)** log₁₀ mean expression of Chr03 single-copy orthogroups. Top row, DE Chr03 (blue) versus FG Chr03 (orange), shown separately for genes inside duplicated blocks (DUP) and in other structural contexts (Other SVs), in rosette and petal. Bottom row, the same values grouped as DUP Chr03 (green) versus Other SVs Chr03 (red), split by subgenome, in rosette and petal. Each pair is annotated with Mann–Whitney U (MW) and Welch’s *t* p-values. DE expression exceeds FG on Chr03 in both tissues and both structural contexts, especially in DUP regions. **(f)** CG methylation metagene profiles on Chr03 for DE (red) versus FG (purple), for genes inside duplicated blocks (left) and outside them (right); axes and per-region Wilcoxon annotation as in **b**. FG hypermethylation upstream of and across gene bodies is most pronounced inside the duplicated blocks.

Despite these methylation asymmetries, transcriptional divergence between the subgenomes was limited (**Fig. 5c**). Differential expression analysis identified only 370 genes in rosette and 222 genes in petal with significant DE-versus-FG expression bias, and expression profiles did not support uniform genome-wide dominance by either subgenome (**Fig. 5c**). This suggests that much of the methylation asymmetry between DE and FG is transcriptionally buffered at the whole-genome scale. A clear exception occurred on Chr03, one of the structurally complex chromosome groups identified above. Single-copy orthogroups on Chr03 showed significantly higher expression from DE than from FG in both petal and rosette tissue (t-test; petal, p = 3.15 × 10^-17^; rosette, p = 1.89 × 10^-16^; **Fig. 5d, Supplementary Table 8**). This signal was unlikely to be explained by read misassignment: an in-silico cross-mapping control showed no positive association between cross-mapping rate and DE-biased expression (Pearson r = −0.27, p < 0.0001, n = 929 gene pairs), and most gene pairs showed no detectable cross-mapping at all (618 of 929 FG genes landing on DE, 672 of 929 DE genes landing on FG; **Supplementary Fig. 2**).

The Chr03 expression bias was closely linked to local structural context. Duplicated sequences showed the strongest association with DE-biased expression (**Fig. 5e**), and these duplicated blocks overlapped the large Chr03 inversion, placing the strongest expression asymmetry within the most structurally complex region of the genome. Methylation differences were also most pronounced in this same region. Single-copy orthogroups within the duplicated and inverted Chr03 blocks showed strong FG hypermethylation in upstream and gene-body regions, whereas the corresponding DE loci were relatively hypomethylated (**Fig. 5f**). Thus, the region with the strongest DE-biased expression also showed the clearest local methylation asymmetry.

To explore whether local subgenome dominance on Chr03 involved genes with shared functional features, we examined biased genes within the duplicated and inverted Chr03 region. Using a threshold of |log₂FC| ≥ 1, we identified 65 DE-biased and 18 FG-biased genes in rosette tissue, and 56 DE-biased and 17 FG-biased genes in petal tissue. The number of GO-annotated genes in this foreground set was too small for robust enrichment analysis (**Supplementary Table 7g–i**). However, qualitative inspection of the strongest DE-biased genes identified *Arabidopsis* orthologues with reported roles in defence signalling and cell-wall remodelling, including SOBIR1 and UGD3. These candidates provide possible functional entry points for future work, but the present data do not establish an adaptive or lineage-specific role for the Chr03 dominance region.

Together, these results show that regulatory divergence between DE and FG is largely buffered genome-wide but becomes locally pronounced on structurally complex Chr03, where duplicated and inverted structure coincides with FG hypermethylation and DE-biased expression.

## Discussion

Here, we presented the first chromosome-level assembly of Alpine Edelweiss, *Leontopodium nivale* subsp*. alpinum*, and used it to resolve the genomic constitution of this culturally iconic alpine plant. The haplotype-resolved assembly shows that Austrian *L. n. alpinum* is a recent allotetraploid composed of two closely related progenitor-derived subgenomes, DE and FG. Comparative analyses link the DE subgenome, among sampled lineages, to Italian *L. n. nivale*, whereas the origin of the FG subgenome remains unresolved. Although the two subgenomes are broadly balanced in gene and repeat content, they differ in localized structural, epigenomic and transcriptional features, culminating in subgenome dominance concentrated on structurally complex Chr03.

The assembly therefore highlights two complementary aspects of Edelweiss genome evolution. First, it provides a framework for resolving the still-incomplete ancestry of the tetraploid. Broader sampling of diploid and polyploid *Leontopodium* lineages across southern Europe and the Asian centre of diversity will be needed to determine whether the sampled Apennine lineage represents the DE progenitor itself or a surviving relative of that lineage, and to identify the missing FG progenitor or its closest extant relative.

Second, Edelweiss may serve as a useful model for studying recent allotetraploidy between closely related genomes. Such systems are especially informative because they capture an early stage of polyploid genome evolution, where progenitor subgenomes remain similar enough to retain broad gene and repeat balance, yet have begun to diverge structurally, epigenomically, and transcriptionally. The localized dominance on Chr03 suggests that early subgenome differentiation need not occur uniformly across the genome, but may instead emerge first in structurally complex regions where recombination, methylation and expression landscapes are perturbed.

Beyond polyploid genome evolution, the assembly provides a foundation for population-level studies of *Leontopodium*. Resequencing across the Alps, Apennines, Balkans, Carpathians, and Asian range of the genus can now be used to reconstruct demographic change, test for selection and identify genomic and epigenomic signatures of adaptation to high-alpine environments. This is particularly relevant under climate change, where altered snow regimes, upward range shifts and habitat contraction may reshape the evolutionary trajectories of alpine plant populations. The genome may also support applied work, including trait mapping, marker-assisted cultivation and breeding strategies for ornamental or medicinal uses, while reducing pressure on wild populations.

In this sense, the Edelweiss genome does more than solve a cytological and phylogenetic puzzle. It begins to reveal the evolutionary history behind one of the most symbolic plants of the European Alps and establishes a foundation for studying how young polyploid genomes stabilize, differentiate, and adapt in high-mountain environments.

## Online Methods

### Plant Sample Collection

Plant material of *Leontopodium nivale* subsp*. alpinum* was collected from single individuals in the Zemmgrund, Zillertal Alps, western Tyrol, Austria, at approximately 2,000 m elevation and approximately 5 km from the Italian border with South Tyrol/Alto Adige (GPS: [47.036758, 11.789298]). Sampling was conducted with permission from the Zillertal Alps Nature Park. For comparative analyses, plant material of *Leontopodium nivale subsp. nivale* was collected from single individuals at Sella di Fonte Fredda, in the Gran Sasso region of the central Apennines, Italy, at ca. 2,000 m elevation (GPS: [42.434267, 13.747583]). Sampling was conducted with permission from the Gran Sasso and Monti della Laga National Park (prot. 0009734/2023). Sampling locations, accession identifiers, tissue types and downstream applications are summarized in Supplementary Table 9.

### PACBIO library preparation, sequencing, and assembly of L. n. alpinum (Austria)

We sequenced *L. n. alpinum* using a whole genome shotgun sequencing strategy and standard sequencing protocols. Sequencing reads were collected using Illumina and PACBIO platforms. Illumina and PACBIO reads were sequenced at the HudsonAlpha Institute in Huntsville, Alabama. Illumina reads were sequenced using the Illumina NovoSeq6000 platform, and the PACBIO reads were sequenced using the REVIO platform using the Circular Consensus Sequencing (CCS) protocol. One 400bp insert 2×150 Illumina fragment library (50.87x coverage) was sequenced along with one 2×150 OmniC library (63.77x coverage) (see Table S1). Prior to assembly, Illumina fragment reads were screened for phix contamination. Reads composed of >95% simple sequence were removed. Illumina reads <50bp after trimming for adapter and quality (q<20) were removed. The final read set consists of 2,573,908,906 reads for a total of 114.64x of high-quality Illumina bases. For the PACBIO sequencing, a total raw sequence yield of 253.0 Gb, with a total coverage of 84.33x (See Table S2).

### Genome assembly and construction of pseudomolecule chromosomes

The version 1.0 assembly was generated by assembling the 14,597,414 PACBIO CCS reads (84.33x coverage) using the HiFiAsm+HIC assembler [30] and subsequently polished using RACON [31]. This produced an initial assembly consisting of 2,415 contigs, with a contig N50 of 21.3 Mb, and a total genome size of 3,404.4 Mb (Table S3).

OmniC Illumina reads from *L. n. alpinum*, were aligned to the initial contig set with bwa mem (Li, 2013), and chromosome scale scaffolding was performed using a combination of the HIC read alignments and the JuiceBox contact map viewer [32]. 92 misjoins were identified in the initial assembly. A total of 262 joins were applied to the assembly to form the final assemblies consisting of 48 chromosomes. Since L. nivale is a tetraploid with no obvious subgenome pairing, chromosomes were grouped in sets of 4 and the set was sorted by size. Subgenomes were denoted as D, E, F, and G. The largest of the group was assigned to “D”, and in descending order to “G”. The resulting sequence was screened for retained vector and/or contaminants. Each chromosome join is padded with 10,000 Ns. Contigs terminating in significant telomeric sequence were identified using the (TTTAGGG)n repeat, and care was taken to make sure that they were properly oriented in the production assembly. The remaining scaffolds were screened against bacterial proteins, organelle sequences, GenBank nr and removed if found to be a contaminant. After forming the chromosomes, it was observed that some small (<20Kb) redundant sequences were present on adjacent contig ends within chromosomes. To resolve this issue, adjacent contig ends were aligned to one another using BLAT [33], and duplicate sequences were collapsed to close the gap between them. A total of 72 adjacent contig pairs were collapsed in the release assembly.

Finally, homozygous SNPs and INDELs were corrected in the release using 53x of Illumina reads (2×150, 400bp insert) by aligning the reads using bwa mem [34] and identifying homozygous SNPs and INDELs with the GATK’s UnifiedGenotyper tool [35]. A total of 1,083 homozygous SNPs and 4,848 homozygous INDELs were corrected in the release. The final version 1.0 release contains 3,147.2 Mb of sequence, consisting of 228 contigs with a contig N50 of 22.0 Mb. A total of 100% of released bases were integrated into chromosomes. The completeness of the genome assembly for *L. n. alpinum* (Austria) was evaluated using BUSCO v5.4.7 [36] with the embryophyta_odb10 lineage dataset.

### DNBSEQ library preparation of L. n. nivale (Italy)

Genomic DNA from *L. n. nivale* from Italy was sequenced by BGI Genomics Co., Ltd. The sequencing involved constructing short-insert libraries using a standard protocol. This included DNA fragmentation, size selection, end repair and “A” tailing, adapter ligation, product size selection, PCR amplification, and library quality control. The libraries were sequenced on the DNBSEQ platform to produce paired-end 150 bp (PE150) reads. The initial raw reads were filtered using SOAPnuke [37] to remove adapter sequences, contaminants, and low-quality reads. The clean data comprised 1.1 Tb for *L. n. nivale* (Italy).

### Genome and transcriptome library preparation and sequencing of L. n. alpinum (Austria)

Genomic DNA from leaf material of *L. n. alpinum* and total RNA from two organs (leaves and bracts) were extracted at the Genome Centre of the Max Planck Institute for Plant Breeding Research and sequenced by BGI Genomics Co., Ltd. using the DNBSEQ platform. A DNBSEQ DNA library was constructed following a standard protocol and sequenced as paired-end 150 bp (PE150) reads, yielding approximately 1,177 million read pairs (353 Gb), with Q20 and Q30 scores above 89% and 77%, respectively, and a GC content of 36%. For the transcriptome, a DNBSEQ eukaryotic strand-specific mRNA library was constructed following a standard protocol. Total RNA was enriched for mRNA using oligo dT beads, fragmented, and then converted into cDNA. The first strand cDNA was synthesized using random primers, and the second strand cDNA was synthesized with dUTP instead of dTTP. The cDNA underwent end-repair, 3’ adenylation, adapter ligation, PCR amplification, quality control, and circularization to form DNA nanoballs (DNBs). Sequencing produced PE150 reads, with approximately 120 million read pairs (30 Gb) per sample, Q20 and Q30 scores above 97% and 93%, respectively, and a GC content of 43%.

### DNA extraction and whole-genome bisulfite sequencing library preparation

DNA was extracted from leaf tissue using Qiagen’s DNeasy Plant Pro Kit (catalog no. 69204, Qiagen) following the manufacturer’s protocol. Briefly, samples in 500 μl Solution CD1 were homogenized using a TissueLyser II (Qiagen) for 3 mins at 25 Hz. DNA was eluted in 50 μl of the supplied elution buffer. Finally, the concentration and purity of DNA samples were assessed using a Qubit dsDNA High Sensitivity (catalog no. Q32854, Invitrogen).

We used NEBNext Ultra II DNA Library Prep Kit (catalog no. E7103, New England BioLabs) for sequencing library preparation and EZ-96 DNA Methylation-Gold MagPrep (catalog no. D5042, ZYMO) for bisulfite treatment. The protocol involved: i) end repair and 3’ adenylation of sonicated DNA fragments by Covaris R230 (Covaris), ii) NEBNext adaptor ligation, iii) cleanup of libraries with AMPure XP Beads (catalog no. A63881, Beckman Coulter), iv) bisulfite treatment, v) PCR enrichment and index ligation using KAPA HiFi Uracil+ Kit (catalog no. KK2802, Agilent) and NEBNext Multiplex Oligos for Enzymatic Methyl-seq (catalog no. E7140L, New England BioLabs) for bisulfite libraries (12 cycles), vi) final cleanup with AMPure XP Beads. Finally, we sequenced paired-end for 150 cycles on a NovaSeq X Plus instrument (Illumina).

### RNA-seq based gene annotation

RNA-seq reads obtained from *L. n. alpinum* (Austria) were aligned with the assembled genome using splice-aware mapper HISAT2 v2.2.1 [39] with default parameters. The aligned reads were assembled and merged with StringTie2 v2.2.2 [40]. GffRead v0.12.7 [41] was used to extract FASTA sequences from the GTF file obtained from StringTie2.

### Prediction and annotation of repeats and transposable elements

We used de-novo TE annotator EDTA [42] to build a repeat library for *L. n. alpinum* (Austria) subgenomes (D, E, F, and G) separately with default parameters. The output included masked subgenomes and their respective repeat libraries. The resulting four masked sub-genomes were merged into a single masked genome file, which was subsequently used by the MAKER v3.01.04 pipeline for genome annotation [43].

### In silico gene model annotation

For gene structure annotation, we employed a combined strategy utilizing both homology-based and transcriptional evidence-based methods. For transcriptional evidence, we integrated the reference transcriptome obtained from StringTie, *L. n. alpinum* transcriptome from the 1000 Plants (1KP) project [44], and mRNA sequences from the entire Gnaphalieae family available on NCBI. These datasets were merged and used as homology-based input evidence for the MAKER pipeline. For protein evidence, we combined the Gnaphalieae protein database from UniProt with *L. n. alpinum* protein data from the 1KP project. Additionally, repeat proteins were identified using the transposable element (TE) protein database supplied by MAKER. For *ab initio* gene prediction, we utilized Exonerate v2.2.0 [45] and AUGUSTUS v3.2.3 [46], both incorporated within the MAKER framework. The annotation process involved two iterative runs of MAKER to refine gene prediction models. After the second iteration, gene models were progressively improved. Non-coding RNAs like tRNAs were predicted by tRNAscan-SE v2.0.12 [47], also run within the MAKER pipeline. The final genome annotation was achieved by merging 48 high-quality gene structure annotation GFF files generated by MAKER, resulting in a comprehensive and accurate annotation of gene structures.

### Structural Variation Analysis

Chromosomal structural variations were identified among the four subgenomes, D, E, F, and G, in *L. n. alpinum* (Austria). To achieve this, each masked homologous chromosome obtained from MAKER was aligned to another counterpart using Minimap2 v2.17 [48]. The alignments were sorted and converted to BAM format using SAMtools v1.19.2 [49]. SyRI v1.6.5 [50] was utilized to do comparative assessments of every chromosome alignment, hence detecting structural variants. Synteny plots were then generated using plotsr v1.1.1 [51] to visualize structural variations among the subgenomes, providing a comprehensive overview of the chromosomal differences between D, E, F, and G subgenomes.

### Genome functional annotation and enrichment

Protein-coding genes predicted from the MAKER pipeline were functionally annotated using the eggNOG-mapper v2.1.12 [52] pipeline and the eggNOG orthology database V5.0.2. For each protein sequence, DIAMOND-based homology searches were performed with default parameters. Gene Ontology (GO) terms were retrieved based on similarities, and a threshold of e-value 0.001 was set. These functional annotations were used for subsequent downstream analyses, including GO enrichment of differentially expressed genes and functional categorization of gene copy number classes.

GO enrichment was performed with topGO v2.58.0 [53] in R v4.4.2, using the eggNOG-mapper GO assignments as the gene-to-GO mapping. Two analyses were run. First, to test whether copy-number reorganization on Chr01 and Chr07 was functionally biased, orthogroups represented in four subgenome copies (retained) and in two copies (depleted) were each tested against a background of all annotated genes on these chromosomes. Second, to characterize the genes within the Chr03 duplicated region, genes falling inside the DUP interval were tested against a background of all annotated genes on Chr03. Enrichment was assessed separately for the biological process, molecular function, and cellular component ontologies.

### Inference of ploidy-type and evolutionary origin

To distinguish between allo- and autotetraploidy, we combined structural variation analysis and gene orthology clustering among four subgenomes. Syntenic block patterns and large structural variations were observed between homoeologous chromosomes for any evidence of clustering. Next, we identified orthologous gene sets (orthogroups) between each homoeologous chromosome set using OrthoFinder v3.0.1 [54]. The input sequences were predicted protein FASTA files obtained from the MAKER pipeline. Orthogroups were identified using DIAMOND-based all-versus-all sequence similarity, followed by gene tree construction and hierarchical orthogroup inference. Finally, we constructed phylogenetic trees for each chromosome set to observe further evidence of clustering. These analyses allowed us to confirm the allotetraploid origin of *L. n. alpinum* (Austria). To establish an upper bound on the mapping rate achievable against this assembly, Austrian *L. n. alpinum* (Austria) short reads were aligned back to the *L. n*. *alpinum* (Austria) reference assembly. Reads were mapped with Minimap2 v2.17 [48] using default parameters, and alignments were filtered to retain primary alignments with MAPQ ≥ 30. The resulting coverage represents the maximum attainable for a conspecific library against this reference and was used as the reference point against which Italian-lineage mapping coverage to Austrian lineage was interpreted. To identify its evolutionary lineage, Minimap2 v2.17 [48] was used to align *L. n. nivale* (Italy) short DNA reads to the *L. n. alpinum* (Austria) whole genome assembly. Alignments were filtered to retain sequences with >MAPQ30. As evidence indicated clustering between DE and FG subgenomes, coverage statistics were averaged across each pair of homoeologous chromosomes (D01-E01, F01-G01, and so on) to identify clusters closely related to *L. n. nivale* (Italy).

### Chloroplast phylogenetics and molecular dating

Chloroplast genomes for the Austrian and Italian *Leontopodium* assemblies were reconstructed from genomic sequencing data using GetOrganelle [55]. A time-calibrated phylogeny was inferred from 96 Asteraceae chloroplast genomes, including 40 *Leontopodium* species, with *Lasiopogon glomerulatus* as the outgroup. Maximum likelihood phylogenetic inference was performed in IQ-TREE 2 [56] with the ModelFinder Plus (MFP) option to identify the optimal nucleotide substitution model, and node support was evaluated using 1,000 ultrafast bootstrap (UFBoot) replicates. Divergence times were estimated using the RelTime Maximum Likelihood method in MEGA11 [57] under the General Time Reversible model with Gamma-distributed rates among sites (GTR+G, 5 discrete rate categories). A partial deletion site-coverage cutoff of 95% was enforced to handle gaps introduced by short-read-derived sequences. To anchor the divergence timeline, two calibration densities were applied in MEGA RelTime, with divergence times obtained from the TimeTree database. Each was specified as a normal distribution with a standard deviation of 0.25 Myr: the *Euchiton involucratus*–*Euchiton japonicus* split (accessions NC_087960.1 and NC_087961.1) was calibrated to a mean of 3.27 Mya (TimeTree median), and the *Helichrysum italicum*–*Pseudognaphalium* split (accessions OL894240.1 and NC_041458.1) to a mean of 6.6 Mya (TimeTree adjusted time, integrating multiple studies). [58].

### Transposable element divergence and insertion age estimation

Gypsy and Copia LTR retrotransposons larger than 1,000 bp were extracted from all four subgenomes and pooled for analysis. Full-precision divergence values (p = 1 − identity) were obtained from RepeatMasker [59] alignment output files using the -a flag, achieving five decimal place resolution. Jukes-Cantor corrected divergence (K) was calculated from raw divergence values, and insertion ages were estimated using the formula T = K / (2r), where r = 1.5 × 10⁻⁸ substitutions per site per year, the standard angiosperm substitution rate. The expected divergence corresponding to the chloroplast point estimate of 20,000 years was calculated as p = μ × t = 1.5 × 10⁻⁸ × 20,000 = 3.0 × 10⁻⁴, and overlaid on the distribution plots together with the 95% confidence interval bounds from the chloroplast dating (CI lower: 801 years, CI upper: 430,600 years).

### Expression analysis of subgenomes

We performed differential expression analysis of DE and FG subgroups in two tissues, rosette and petal leaves, using DESeq2 [60]. Counts from D and E homoeologues were grouped as the DE subgenome, and counts from F and G homoeologues as the FG subgenome. Subgroups DE and FG were assigned as control and treatment, respectively. This was done for both tissues separately. For consistency, we used single-copy orthologs to quantify their gene expression patterns. A threshold of adjusted p-value < 0.05 and absolute log2 fold change > 1 was used to obtain significant DEGs. Afterwards, plots of heatmaps, volcano plots, and log2fold change for both tissues were generated. In order to understand the overall expression levels between the DE and FG subgenomes, we quantified chromosome-wise average normalized expression levels in upregulated genes and generated bar plots for each tissue.

To test whether DE subgenome expression dominance on Chr03 could be attributable to cross-mapping of reads between homeologous gene copies, we performed an in-silico simulation control. Synthetic 150 bp paired-end reads were generated separately from DE and FG Chr03 transcript sequences (1,282 single-copy orthologous gene pairs) using wgsim (∼500 read pairs per transcript), providing read sets of known subgenome origin. Reads were mapped to the full tetraploid reference using identical HISAT2 [30] parameters to the original expression analysis, and gene-level counts were generated using identical featureCounts [61] settings. The cross-mapping rate was calculated per gene pair as the proportion of reads simulated from one subgenome that were assigned to the homeologous gene on the other subgenome. The relationship between cross-mapping rate and the real DE-versus-FG Wald statistic from the DESeq2 [60] expression analysis was assessed using Pearson correlation.

To further understand the functional roles of DE and FG subgroups, we wrote in-house scripts to perform functional enrichment for upregulated genes in DE and FG subgroups for each tissue. Raw gene expression counts were compared between the DE and FG subgenomes to quantify expression differences between homoeologous gene pairs. For each orthologous pair, a t-statistic was calculated to assess the magnitude and direction of expression divergence between subgenomes. To evaluate the relationship between genome structure and expression bias, t-statistics were correlated with structural variation categories across the genome. Expression differences of orthologous genes located within duplicated regions were further compared with those associated with other types of structural variation using custom in-house scripts. Statistical significance of expression differences between groups was assessed using two-sided Mann–Whitney U tests.

### DNA methylome analysis

In order to support evidence of subgenome functionalization observed from expression analysis, we wanted to see whether it is reflected in the methylome profile of DE vs. FG subgenomes. The WGBS reads were mapped to the *L. n. alpinum* (Austria) genome and analyzed using MethylStar pipeline v1.0, which employs METHimpute for cytosine-level methylation calling [62]. For gene and TE, methylation metaplots were generated for DE and FG subgenomes in CG, CHG, and CHH contexts. To support expression analyses, methylation profiles for significant DEGs in each tissue were plotted for DE vs. FG subgenomes. All plots were made in three bins, with ±2 kb flanking regions, transcription start site (TSS), and transcription end site (TES).

## Supplemental Information

Supplementary Table 1. Details on chromosome sizes, assembly statistics, and completeness.

Supplementary Table 2. Genome sizes vs. repeat content correlation among 23 plant species.

Supplementary Table 3. Genome-wide and per-chromosome wise orthogroups, and orthologs copy-number distribution.

Supplementary Table 4. Alignment statistics of *L. n. alpinum* (Austria) short-read to *L. n. alpinum* (Austria) assembly, and *L. n. nivale* (Italy) to *L. n. alpinum* (Austria) assembly. SNP calls for against each subgenome.

Supplementary Table 5. Chloroplast genome accessions used for time-calibrated phylogenetic analysis 102 Asteraceae taxa.

Supplementary Table 6. Structural Variation summaries for comparisons between D and E; E and F; F and G.

Supplementary Table 7. GO term enrichment among Chr01+Chr07, and in DUP region of Chr03.

Supplementary Table 8. Mean expression differences t-test for each homoeologous chromosome between DE and FG subgenomes.

Supplementary Table 9. Plant sample data.

## Acknowledgements

We thank Willi Seifert from the Zillertal High-alps Nature Park and Carlo Catonica from the Gran Sasso and Monti della Laga National Park for their cooperation and permission to collect plant material. We thank members of the Klaus Mayer group for helpful discussion, for help with sample collection in pilot studies. FJ was partly supported by grant ATR2025-164499 funded by MICIU/AEI /10.13039/501100011033. This work was also supported by grants CEX2019-000902-S and CEX2025-001593-S funded by MICIU/AEI/10.13039/501100011033, and by the CERCA Programme / Generalitat de Catalunya.

## Supplementary figures

**Supplementary Fig. 1.**
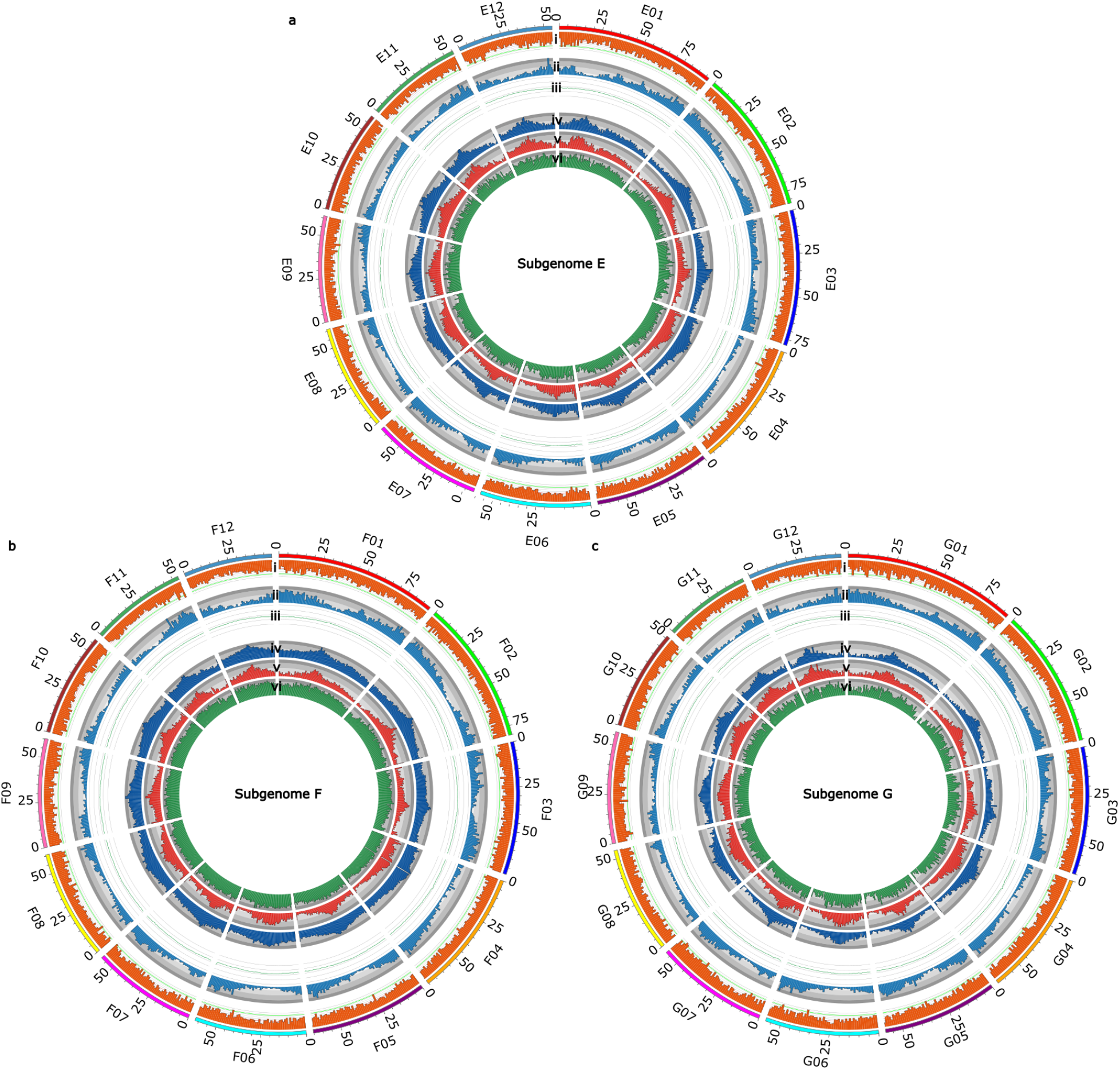
A haplotype-resolved, chromosome-scale assembly resolves four structurally balanced chromosome sets in *Leontopodium nivale* subsp*. alpinum.* Circos plot of subgenome (a) E, (b) F and, (c) G showing six tracks in 1 Mb bins: (i) TE density (∼65–67%), (ii) gene density, (iii) GC content (∼36%), (iv) CG methylation, (v) CHG methylation, and (vi) CHH methylation.

**Supplementary Fig. 2.**
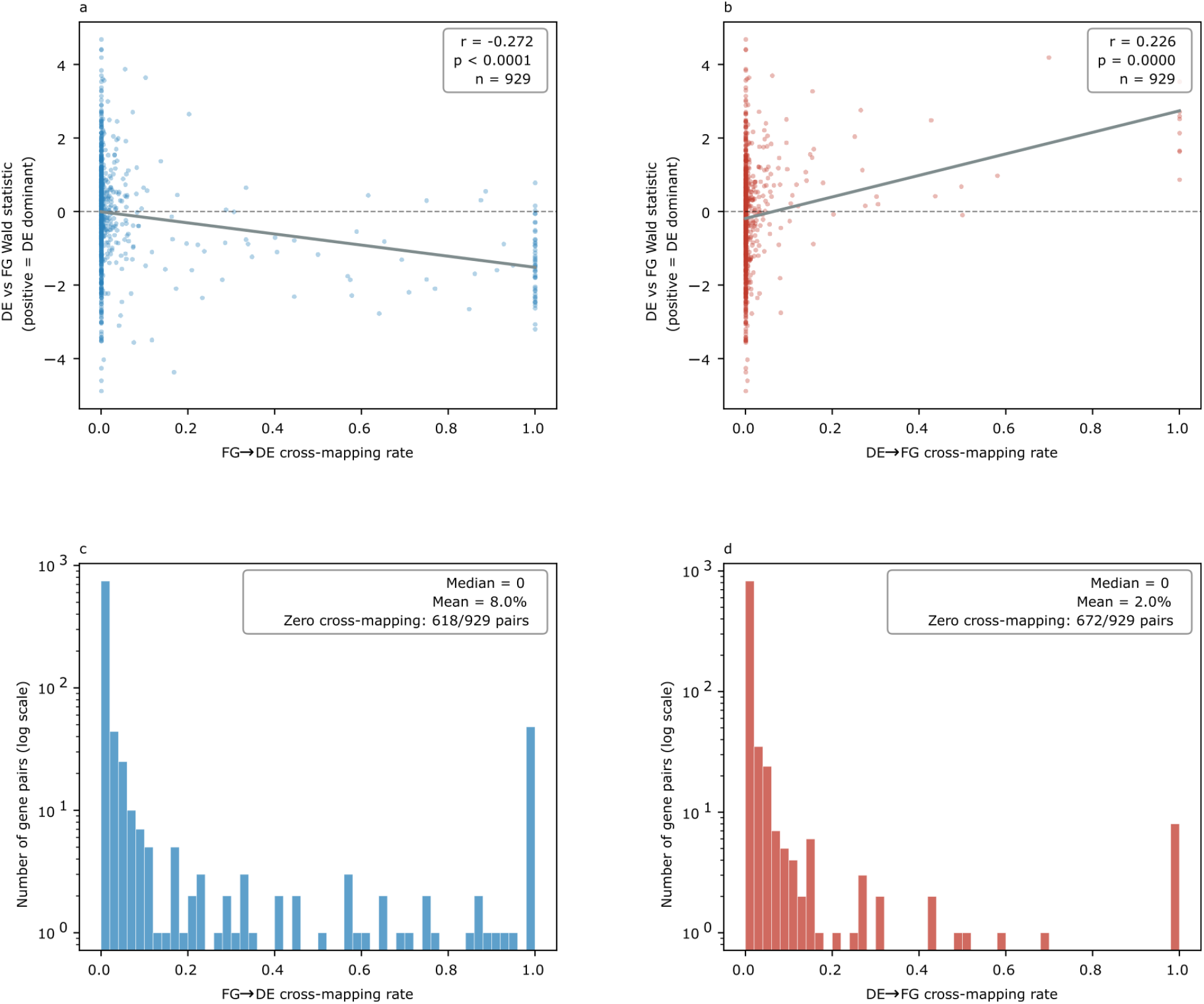
In-silico cross-mapping control for Chr03 DE-biased expression. Test of whether the DE-biased expression of Chr03 single-copy orthogene pairs could arise from sequencing reads misassigning between the homoeologous DE and FG copies rather than from genuine subgenome expression bias. For each of 929 DE–FG orthogene pairs on Chr03, the cross-mapping rate, the fraction of one copy’s reads that align to the opposite copy — was quantified in each direction [in-silico method to confirm] and related to that pair’s expression bias, summarised by the DE-versus-FG Wald statistic (positive values indicate DE-dominant expression). **(a)** Wald statistic versus FG→DE cross-mapping rate (FG reads aligning to the DE copy). Were DE dominance an artefact of FG reads leaking onto DE, this association would be positive; instead it is weakly negative (Pearson r = −0.272, P < 0.0001, n = 929), opposite to the direction expected from misassignment. Solid line, linear fit; dashed line, Wald = 0. **(b)** Wald statistic versus DE→FG cross-mapping rate (Pearson r = 0.226, P < 0.0001, n = 929). **(c)** Distribution of FG→DE cross-mapping rate across the 929 pairs (y, log scale). Cross-mapping is absent for the majority of pairs (median 0; zero cross-mapping in 618 of 929 pairs; mean 8.0%). **(d)** Distribution of DE→FG cross-mapping rate (median 0; zero cross-mapping in 672 of 929 pairs; mean 2.0%).

## References

1. Ischer M, Dubuis A, Keller R, Vittoz P. A better understanding of the ecological conditions for Leontopodium alpinum Cassini in the Swiss Alps. Folia Geobot. 2014;49: 541–558.

2. Von Wildenbruch E, Stille JS, Wissemann V. Das Edelweiß: (K)ein Europäer!

3. Vigneron JP. Optical structure and function of the white filamentary hair covering the edelweiss bracts. Phys Rev E. 2005;71: 011906.

4. Safer S, Tremetsberger K, Guo Y-P, Kohl G, Samuel MR, Stuessy TF, et al. Phylogenetic relationships in the genus Leontopodium (Asteraceae: Gnaphalieae) based on AFLP data: PHYLOGENETIC RELATIONSHIPS IN LEONTOPODIUM. Bot J Linn Soc. 2011;165: 364–377.

5. Blöch C, Dickoré WB, Samuel R, Stuessy TF. MOLECULAR PHYLOGENY OF THE EDELWEISS (LEONTOPODIUM, ASTERACEAE – GNAPHALIEAE). Edinburgh J Bot. 2010;67: 235–264.

6. Xu X-M, Wei Z, Sun J-Z, Zhao Q-F, Lu Y, Wang Z-L, et al. Phylogeny of Leontopodium (Asteraceae) in China—with a reference to plastid genome and nuclear ribosomal DNA. Front Plant Sci. 2023;14: 1163065.

7. Greuter, W. 2006+: Compositae (pro parte majore). – In: Greuter, W. & Raab-Straube, E. von, Compositae. Euro+Med PlantBase - the information resource for Euro-Mediterranean plant diversity. Published at https://europlusmed.org/.

8. Stille JS, Jaeger M, Dickoré WB, Ehlers K, Holzhauer SIJ, Mayland-Quellhorst E, et al. CHROMOSOME NUMBERS OF THE EDELWEISS, LEONTOPODIUM (ASTERACEAE, COMPOSITAE – GNAPHALIEAE). Edinb J Bot. 2014;71: 23–33.

9. Van de Peer Y, Mizrachi E, Marchal K. The evolutionary significance of polyploidy. Nat Rev Genet. 2017;18: 411–424.

10. Ramsey J, Schemske DW. Pathways, mechanisms, and rates of polyploid formation in flowering plants. Annu Rev Ecol Syst. 1998;29: 467–501.

11. Leitch AR, Leitch IJ. Genomic plasticity and the diversity of polyploid plants. Science. 2008;320: 481–483.

12. McClintock B. The significance of responses of the genome to challenge. Science. 1984;226: 792–801.

13. Bottani S, Zabet NR, Wendel JF, Veitia RA. Gene expression dominance in allopolyploids: Hypotheses and models. Trends Plant Sci. 2018;23: 393–402.

14. Chen ZJ. Genetic and epigenetic mechanisms for gene expression and phenotypic variation in plant polyploids. Annu Rev Plant Biol. 2007;58: 377–406.

15. Song Q, Chen ZJ. Epigenetic and developmental regulation in plant polyploids. Curr Opin Plant Biol. 2015;24: 101–109.

16. Edger PP, Smith R, McKain MR, Cooley AM, Vallejo-Marin M, Yuan Y, et al. Subgenome Dominance in an Interspecific Hybrid, Synthetic Allopolyploid, and a 140-Year-Old Naturally Established Neo-Allopolyploid Monkeyflower. Plant Cell. 2017;29: 2150–2167.

17. Buggs RJA, Elliott NM, Zhang L, Koh J, Viccini LF, Soltis DE, et al. Tissue-specific silencing of homoeologs in natural populations of the recent allopolyploid Tragopogon mirus. New Phytol. 2010;186: 175–183.

18. Kokot M, Dlugosz M, Deorowicz S. KMC 3: counting and manipulating k-mer statistics. Bioinformatics. 2017;33: 2759–2761.

19. Ranallo-Benavidez TR, Jaron KS, Schatz MC. GenomeScope 2.0 and Smudgeplot for reference-free profiling of polyploid genomes. Nat Commun. 2020;11: 1432.

20. Parisod C, Salmon A, Zerjal T, Tenaillon M, Grandbastien M-A, Ainouche M. Rapid structural and epigenetic reorganization near transposable elements in hybrid and allopolyploid genomes in Spartina. New Phytol. 2009;184: 1003–1015.

21. Petit M, Guidat C, Daniel J, Denis E, Montoriol E, Bui QT, et al. Mobilization of retrotransposons in synthetic allotetraploid tobacco. New Phytol. 2010;186: 135–147.

22. Vicient CM, Casacuberta JM. Impact of transposable elements on polyploid plant genomes. Ann Bot. 2017;120: 195–207.

23. Baduel P, Quadrana L, Hunter B, Bomblies K, Colot V. Relaxed purifying selection in autopolyploids drives transposable element over-accumulation which provides variants for local adaptation. Nat Commun. 2019;10: 5818.

24. Mason AS, Wendel JF. Homoeologous Exchanges, Segmental Allopolyploidy, and Polyploid Genome Evolution. Front Genet. 2020;11: 1014.

25. Zhao J, Li J, Lv R, Wang B, Zhang Z, Yu T, et al. Meiotic pairing irregularity and homoeologous chromosome compensation cause rapid karyotype variation in synthetic allotetraploid wheat. New Phytol. 2023;239: 606–623.

26. Huang K, Rieseberg LH. Frequency, Origins, and Evolutionary Role of Chromosomal Inversions in Plants. Front Plant Sci. 2020;11: 296.

27. Zhang X, Hu Y, Huang K, Wright SI, Rieseberg LH. Recombination suppression in plant adaptation and speciation. New Phytologist. 2026;250: 2061–2077.

28. Freeling M. Bias in plant gene content following different sorts of duplication: tandem, whole-genome, segmental, or by transposition. Annu Rev Plant Biol. 2009;60: 433–453.

29. Tasdighian S, Van Bel M, Li Z, Van de Peer Y, Carretero-Paulet L, Maere S. Reciprocally Retained Genes in the Angiosperm Lineage Show the Hallmarks of Dosage Balance Sensitivity. Plant Cell. 2017;29: 2766–2785.

30. Cheng H, Concepcion GT, Feng X, Zhang H, Li H. Haplotype-resolved de novo assembly using phased assembly graphs with hifiasm. Nat Methods. 2021;18: 170–175.

31. Vaser R, Sović I, Nagarajan N, Šikić M. Fast and accurate de novo genome assembly from long uncorrected reads. Genome Res. 2017;27: 737–746.

32. Robinson JT, Turner D, Durand NC, Thorvaldsdóttir H, Mesirov JP, Aiden EL. Juicebox.js Provides a Cloud-Based Visualization System for Hi-C Data. Cell Syst. 2018;6: 256–258.e1.

33. Kent WJ. BLAT--the BLAST-like alignment tool. Genome Res. 2002;12: 656–664.

34. Li H. Aligning sequence reads, clone sequences and assembly contigs with BWA-MEM. arXiv [q-bio.GN]. 2013. doi:10.48550/arXiv.1303.3997

35. McKenna A, Hanna M, Banks E, Sivachenko A, Cibulskis K, Kernytsky A, et al. The Genome Analysis Toolkit: a MapReduce framework for analyzing next-generation DNA sequencing data. Genome Res. 2010;20: 1297–1303.

36. Manni M, Berkeley MR, Seppey M, Zdobnov EM. BUSCO: Assessing Genomic Data Quality and Beyond. Curr Protoc. 2021;1: e323.

37. Chen Y, Chen Y, Shi C, Huang Z, Zhang Y, Li S, et al. SOAPnuke: a MapReduce acceleration-supported software for integrated quality control and preprocessing of high-throughput sequencing data. Gigascience. 2018;7: 1–6.

38. Hu T, Chen J, Lin X, He W, Liang H, Wang M, et al. Comparison of the DNBSEQ platform and Illumina HiSeq 2000 for bacterial genome assembly. Sci Rep. 2024;14: 1292.

39. Kim D, Paggi JM, Park C, Bennett C, Salzberg SL. Graph-based genome alignment and genotyping with HISAT2 and HISAT-genotype. Nat Biotechnol. 2019;37: 907–915.

40. Kovaka S, Zimin AV, Pertea GM, Razaghi R, Salzberg SL, Pertea M. Transcriptome assembly from long-read RNA-seq alignments with StringTie2. Genome Biol. 2019;20: 278.

41. Pertea G, Pertea M. GFF utilities: GffRead and GffCompare. F1000Res. 2020;9: 304.

42. Ou S, Su W, Liao Y, Chougule K, Agda JRA, Hellinga AJ, et al. Benchmarking transposable element annotation methods for creation of a streamlined, comprehensive pipeline. Genome Biol. 2019;20: 275.

43. Cantarel BL, Korf I, Robb SMC, Parra G, Ross E, Moore B, et al. MAKER: an easy-to-use annotation pipeline designed for emerging model organism genomes. Genome Res. 2008;18: 188–196.

44. One Thousand Plant Transcriptomes Initiative. One thousand plant transcriptomes and the phylogenomics of green plants. Nature. 2019;574: 679–685.

45. Slater GSC, Birney E. Automated generation of heuristics for biological sequence comparison. BMC Bioinformatics. 2005;6: 31.

46. Stanke M, Keller O, Gunduz I, Hayes A, Waack S, Morgenstern B. AUGUSTUS: ab initio prediction of alternative transcripts. Nucleic Acids Res. 2006;34: W435–9.

47. Chan PP, Lin BY, Mak AJ, Lowe TM. tRNAscan-SE 2.0: improved detection and functional classification of transfer RNA genes. Nucleic Acids Res. 2021;49: 9077–9096.

48. Li H. Minimap2: pairwise alignment for nucleotide sequences. Bioinformatics. 2018;34: 3094–3100.

49. Li H, Handsaker B, Wysoker A, Fennell T, Ruan J, Homer N, et al. The Sequence Alignment/Map format and SAMtools. Bioinformatics. 2009;25: 2078–2079.

50. Goel M, Sun H, Jiao W-B, Schneeberger K. SyRI: finding genomic rearrangements and local sequence differences from whole-genome assemblies. Genome Biol. 2019;20: 277.

51. Goel M, Schneeberger K. Plotsr: Visualizing structural similarities and rearrangements between multiple genomes. Bioinformatics. 2022;38: 2922–2926.

52. Cantalapiedra CP, Hernández-Plaza A, Letunic I, Bork P, Huerta-Cepas J. EggNOG-mapper v2: Functional annotation, orthology assignments, and domain prediction at the metagenomic scale. Mol Biol Evol. 2021;38: 5825–5829.

53. topGO: Enrichment Analysis for Gene Ontology. In: R-universe [Internet]. [cited 14 July 2026]. Available: https://bioc.r-universe.dev/topGO

54. Emms DM, Kelly S. OrthoFinder: phylogenetic orthology inference for comparative genomics. Genome Biol. 2019;20: 238.

55. Jin J-J, Yu W-B, Yang J-B, Song Y, dePamphilis CW, Yi T-S, et al. GetOrganelle: a fast and versatile toolkit for accurate de novo assembly of organelle genomes. Genome Biol. 2020;21: 241.

56. Minh BQ, Schmidt HA, Chernomor O, Schrempf D, Woodhams MD, von Haeseler A, et al. IQ-TREE 2: New Models and Efficient Methods for Phylogenetic Inference in the Genomic Era. Mol Biol Evol. 2020;37: 1530–1534.

57. Tamura K, Stecher G, Kumar S. MEGA11: Molecular Evolutionary Genetics Analysis Version 11. Mol Biol Evol. 2021;38: 3022–3027.

58. Kumar S, Suleski M, Craig JM, Kasprowicz AE, Sanderford M, Li M, et al. TimeTree 5: An Expanded Resource for Species Divergence Times. Mol Biol Evol. 2022;39. doi:10.1093/molbev/msac174

59. Website. Available: Smit, A. F. A., Hubley, R. & Green, P. RepeatMasker Open-4.0. http://www.repeatmasker.org (2013–2015).

60. Love MI, Huber W, Anders S. Moderated estimation of fold change and dispersion for RNA-seq data with DESeq2. Genome Biol. 2014;15: 550.

61. Liao Y, Smyth GK, Shi W. featureCounts: an efficient general purpose program for assigning sequence reads to genomic features. Bioinformatics. 2014;30: 923–930.

62. Shahryary Y, Hazarika RR, Johannes F. MethylStar: A fast and robust pre-processing pipeline for bulk or single-cell whole-genome bisulfite sequencing data. BMC Genomics. 2020;21: 479.

